# Inflammation-mediated induction of Clec18a inhibits immune responses via Trpm7-dependent Cbl activation

**DOI:** 10.64898/2026.09.24.754070

**Authors:** Zhengyu Jiang, Lulong Bo, Na Li, Huabiao Chen, Tao Li, Yan Zhang, Xiaoming Deng, Jinjun Bian

**Author notes:** These authors contributed equally (Zhengyu Jiang, Lulong Bo, Na Li, Huabiao Chen). Correspondence (J. B.), (X. D.), (Y. Z.).

## Abstract

Innate immune responses must be tightly controlled to maintain immuno-homeostasis. However, the regulatory mechanisms underlying the activation and early stages of the innate immune response remain poorly defined. Here, we found the soluble C-type lectin Clec18a is up-regulated in multiple inflammatory diseases, including sepsis and influenza, and secreted within minutes upon activation in multiple immune cells, including macrophages, dendritic cells (DCs), and T cells. Clec18a-myeloid-knockout mice exhibited reduced survival and exacerbated inflammation than the wild-type. However, exogenous recombinant Clec18a administration alleviated inflammation both *in vitro* and *in vivo*. Mechanistically, we identified macrophage-inducible C-type lectin (Mincle) as the receptor of Clec18a. Clec18a-Mincle binding triggers Trpm7-dependent calcium influx, which activates the ubiquitin E3 ligase Cbl and leads to Mincle-Cbl-MyD88-Trpm7 complex formation and degradation. Cbl deficiency, calcium inhibition and Trpm7 knockdown abrogated Clec18a-mediated immune suppression. Thus, our study revealed Clec18a as a critical checkpoint on the initiation of innate immune response through an intrinsic crosstalk between CLR and TLR signaling.

**Graphical abstract:** Inflammation-induced Clec18a secretion binds to Mincle, triggering Trpm7-mediated extracellular calcium influx, directly activates Cbl and orchestrates the assembly of the Mincle-Trpm7-Cbl-MyD88 complex. Subsequent ubiquitination leads to proteasomal degradation of this complex, thereby abrogating CLR-TLR signaling crosstalk.

## INTRODUCTION

Pattern-recognition receptors (PRRs), such as toll-like receptors (TLRs) and C-type lectin receptors (CLRs), initiate inflammatory responses upon activation by pathogen-associated molecular patterns (PAMPs) and damage-associated molecular patterns (DAMPs).[1] The binding of PAMPs and DAMPs to TLRs triggers myeloid differentiation primary response protein 88 (MyD88)-dependent or MyD88-independent signaling, resulting in the activation of NF-κB and the mitogen-activated protein kinase (MAPK) signaling pathway and the production of inflammatory cytokines.[1, 2, 3] Appropriate cytokine production helps establish protective immunity and promotes pathogen clearance, whereas uncontrolled inflammation can result in organ injury, immune disorders, and even sepsis or autoimmune diseases.[1, 4]

Multiple negative autoregulatory loops have evolved to prevent immune hyperactivity.[1, 5] Generally, the intracellular regulation initiated either concurrently with or after the activation of the signaling pathway, induing transcriptional, translational and posttranslational modifications, as well as epigenetic and metabolic reprogramming.[3, 6, 7, 8, 9] In contrast, secretory molecules, such as IL-10 and TGF-β, exert systematic regulatory effects by participating in autocrine and/or paracrine signaling, typically produced during the late phase of inflammation, facilitating inflammation resolution and tissue repair.[10, 11, 12] However, it remains unclear how immune cells “tune” the responses and the role of secretory molecule in the ultra-early stage during inflammation.

C-type lectins include transmembrane or soluble proteins that mediate immune responses to pathogens and cell death, contributing to the maintenance of immune homeostasis.[13, 14] Macrophage-inducible C-type lectin (Mincle), a transmembrane C-type lectin receptor (CLR) of the Dectin-2 group, is activated when a ligand binds and signal through Fc receptor γ-chain (FcRγ) chain, leading to the activation of the Syk-PKCδ-PLCγ axis and NF-κB via the Card9-Bcl10-Malt1 complex.[15, 16] CLRs have been shown to interact with TLRs via multiple pathways.[17] For example, activation of the DC immunoreceptor (DCIR) recruits SHP-1/2 to inhibit TLR- and MyD88-dependent signaling and cytokine production.[18, 19] Moreover, blood DC antigen 2 (BDCA2) inhibits MyD88 via calcineurin-mediated tonic calcium signaling.[20, 21] However, these findings are based on dual stimulation of CLR and TLR signaling simultaneously, which necessitates concurrent PAMP binding; it remains unclear whether there is intrinsic crosstalk between TLR and CLR signaling.

C-type lectin 18A (Clec18a), a soluble C-type lectin in the CLR family, is abundantly expressed in peripheral blood cells, as well as in the liver, kidney and heart.[22] In human monocytes, Clec18a is upregulated during differentiation into macrophages or dendritic cells.[23] Previous studies have reported that Clec18a can enhance the host immune response to viral infection and may serve as a biomarker to predict the outcomes of patients with Hepatitis C virus (HCV) infection.[22, 24] In macrophages, Clec18a can impair phagocytosis by inhibiting FcRγ activation.[25] However, the expression, specific receptors, and detailed mechanism of Clec18a in the innate immune response require further investigation.

In the present study, we demonstrated that Clec18 functions not only as an early “responder” to diverse stimuli in multiple cells but also as a “suppressor” of inflammatory responses both *in vitro* and *in vivo*. Moreover, we identified Mincle as the receptor of Clec18a. The binding of Clec18a to Mincle induces Trpm7-dependent extracellular calcium influx, which activates the E3 ubiquitin ligase Cbl and subsequently promotes Mincle-Cbl-MyD88 complex formation and degradation. Our study reveals the novel role of Clec18a as an intrinsic negative regulator of the acute immune response and provides new insights into the intrinsic crosstalk between CLRs and TLRs in the acute phase of the immune response.

## RESULTS

### Inflammation induces rapid Clec18a production

To explore the potential involvement of Clec18a in inflammatory responses, we first analyzed its transcriptional level across a variety of inflammatory conditions using publicly available datasets from the Gene Expression Omnibus (GEO). As shown in Figure 1A, Clec18a expression was significantly elevated (logFC > 0) in the blood of patients with sepsis, bacterial pneumonia, influenza, severe community-acquired pneumonia and autoimmune disease like psoriasis (normalized Clec18a expression in Table S1). These results suggested a potential role of Clec18a in inflammatory settings. Given that sepsis represents a classical model for innate immune responses, we selected sepsis as the primary experimental model to dissect the functional role of Clec18a in inflammation. We demonstrated Clec18a was upregulated in merged sepsis datasets of whole blood transcription (Figure 1B). Moreover, Gene Set Enrichment Analysis (GSEA) of the merged data showed that samples with high Clec18a expression exhibited significant enrichment of key inflammatory pathways (Figure S1A), suggesting the potential role during inflammation. In our own clinical cohort of patients with postoperative abdominal or bloodstream infections (demographic in Table S2), conditions that partially recapitulate sepsis, we also observed elevated Clec18a levels in both PBMC-derived mRNA and serum samples (Figure 1C and 1D), further supporting its association with inflammatory states.

**Figure 1.**
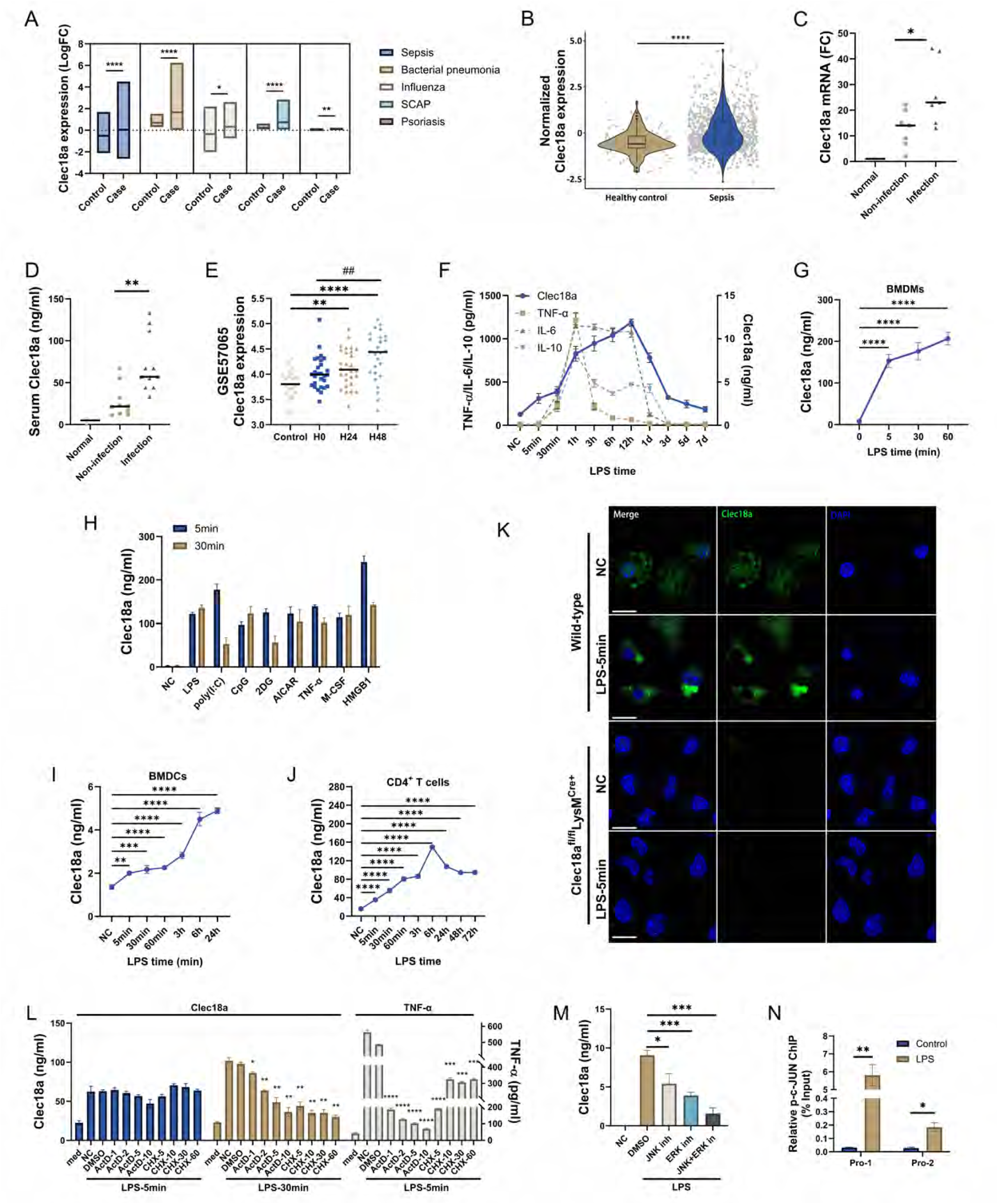
Inflammation induces rapid Clec18a production. A: Fold-change of Clec18a in datasets of GEO, sepsis (GSE137340, GSE26440, GSE57065 and GSE65682, case n=959, control n=111), Bacteria pneumonia (GSE20346, case n=26, control n=18), Influenza (GSE20346, case n=19, control n=18), severe community acquired pneumonia (SCAP, GSE196399, case n=56, control n=21), Psoriasis (GSE147339, case n=10, control n=10). B: Normalized transcriptional level of Clec18a in merged datasets of blood samples from sepsis patients and healthy controls in GEO database (n=1070). C, D: Fold-change of Clec18a in PBMCs (C) and serum Clec18a level (D) of patients in intensive care units with infection (n=7 for mRNA in PBMCs and n=11 in serum Clec18a analysis), patients without infection (n=7 for mRNA in PBMCs and n=10 in serum Clec18a analysis), and healthy volunteers (n=3). E: Normalized transcriptional level of Clec18a of different timepoints of sepsis patients in GSE57065 (n=107), (Blood samples collected within 30 minutes (H0) (n=28), 24 (H24) (n=28) and 48 (H48) (n=26) hours after sepsis diagnosis, healthy control n=25). F: Cytokine and Clec18a levels in the serum of C57BL/6 mice injected with LPS *i.v.* (50 μg/g) at various time points (n=5 per group). G: Levels of Clec18a in the supernatant of bone marrow-derived macrophages (BMDMs) from mice treated with lipopolysaccharide (LPS) (100 ng/ml) at various time points (n=3). H: Levels of Clec18a in the supernatant of BMDMs treated with various stimulators and time points (LPS, 100 ng/ml; poly(I:C), 5 μg/ml; CpG, 0.2 μM; 2-DG, AICAR, 500 μM; TNF-α, 50 ng/ml; M-CSF, 25 ng/ml; HMGB1, 5 μg/ml) (n=5). Data of each stimuli in five and thirty minutes were all significantly different (*p*<0.001 or *p*<0.0001) from NC group. I, J: Levels of Clec18a in the supernatant of mouse CD11c^+^ BMDCs (I), human CD4^+^ T cells (J), and stimulated with LPS (10 ng/ml) or anti-CD3 and anti-CD28 antibodies, respectively, at various time points (n=4). K: Confocal microscopy images of Clec18a expression in BMDMs from Clec18a^fl/fl^LysM^Cre+^ and littermate wild-type mice stimulated with LPS at 5 minutes (Bar=5 μM). L: Levels of Clec18a and TNF-α in the supernatant of BMDMs treated with indicated inhibitors at different concentrations (Act D at 1, 2, 5, 10 μM; CHX at 5, 10, 30, 60 μM, *compared to the DMSO group) for 30 min and then stimulated with LPS for 5 or 30 min (n=3). M: Levels of Clec18a in the supernatant of BMDMs pre-treated with the indicated inhibitors (Erk1/2 inhibitor 10 μM, Jnk inhibitor (SP600125, 10 μM); p38 inhibitor (SB203580, 10 μM); p65 inhibitor (SC75741, 10 μM)) for 30min and then stimulated with LPS for six hours (n=3). N: Relative expression of the promoter region of Clec18a immunoprecipitated with phospho-c-Jun (n=3). *p<0.05, ***p<0.001, ****p<0.0001; Student’s *t* test (A, B), one-way ANOVA (C, D, E*, G, I, J, L, M), repeated measures ANOVA (E^#^); two-way ANOVA (N); n=biological replicates for in vivo assay and replicate wells for in vitro assay; Data shown in all panels are representative of at least three independent experiments.

Inflammation contains multiphasic dynamics. Indeed, Clec18a transcription quickly up-regulated within 24 hours and continued elevating in sepsis (Figure 1E). To characterize the kinetic expression of Clec18a during inflammation, we analyzed serum Clec18a level *in vivo* of mice in responses to several TLR agonists. In lipopolysaccharide-(LPS) challenged (*i.v.*) mice, Clec18a was quickly elevated in serum at five minutes post-challenge, earlier than the rises of TNF-α, IL-6 and the classic anti-inflammatory cytokine IL-10. It also maintained elevation as inflammatory progression and declined only after the resolution of TNF-α, IL-6 and IL-10 (Figure 1F, Figure S1B). This kinetic feature was also observed in CpG- and poly(I:C)-challenged mice, that Clec18a increased earlier than canonical cytokines (Figure S1C and S1D). These results indicated Clec18a as an early-responder during inflammation.

We then investigated the Clec18a production in macrophage, a major source of cytokines during inflammations. We stimulated murine bone marrow-derived macrophages (BMDMs) with LPS. Notably, Clec18a expression increased within 5 minutes of LPS challenge, reaching ∼70% of the peak level that was achieved at 1 hour (Figure 1G). In addition to TLR, diverse stimuli, including proinflammatory cytokines (TNF-α, M-CSF), metabolic stressors (2-DG, AICAR), and the DAMP molecule HMGB1, elicited comparable increases in Clec18a levels within 5 minutes of challenge (Figure 1H) in macrophage. In addition to innate immune cells, bone marrow-derived dendritic cells (BMDCs) and CD4^+^ T cells activated with LPS or CD3/CD28 engagement, respectively, also showed rapid Clec18a production (Figure 1I and 1J). In human immune cells, LPS-stimulated CD14^+^ monocytes from PBMCs and PMA-differentiated THP-1 macrophages showed similarly rapid Clec18a upregulation (Figure S1E and S1F). Taken together, these data indicate that Clec18a rapidly responds to various stimuli in both innate and adaptive immune cells.

The rapid production of Clec18a prompted us to investigate its storage in cells. Immunoblotting (Figure S1G) revealed rapid increases in Clec18a expression five minutes post-LPS challenge. To visualize the basal storage of Clec18a, we generated myeloid-specific Clec18a knockout mice (Clec18afl/flLysMCre+) and analyzed Clec18a in unstimulated and five minutes post-LPS stimulation in BMDMs through confocal microscopy. Notably, compared to Clec18a-deficient macrophage, wild-type macrophage contains relative Clec18a expression in unstimulated macrophage, mainly distributed in vesicle-like compartment and quickly enhanced upon stimulation in five minutes (Figure 1K). Moreover, this rapid increase was not abrogated after treatment with actinomycin D (Act D) or cycloheximide, CHX, contrasting to the TNF-*α*, a classic de novo-synthesized cytokine (Figure 1L), but abrogated with treatment of Brefeldin A, an inhibitor of anterograde transport from the endoplasmic reticulum to the Golgi apparatus that inhibits protein secretion (Figure S1H). However, Act D and CHX could decrease Clec18a expression in the late stage of the immune response (30 minutes post-LPS) (Figure 1L). These data suggested that LPS increased the basal production of Clec18a in the early stage of the immune response and Clec18a synthesis took place in the late stage of the immune response. We then investigated the transcriptional mechanism of Clec18a 30 minutes after LPS stimulation. After TLR4 recognizes and binds to stimuli, it initiates the activation of the JNK/ERK/p38/p65 signaling pathway.[2] We first pharmacologically inhibited the members of this pathway to analyze those required for Clec18a induction. The results revealed that inhibiting JNK and ERK attenuated Clec18a expression, with the combined blockade resulting in additive suppression (Figure 1M). Chromatin immunoprecipitation confirmed the involvement of the AP-1 transcription factor c-Jun, a known ERK/JNK target, in Clec18a transcriptional regulation (Figure 1N). These results suggested that stored Clec18a was the source of Clec18a in the early stage of inflammation, whereas Clec18a is produced via de novo synthesis, which is dependent on AP-1 regulation, in the late stage of inflammation.

Collectively, our data demonstrated that Clec18a is rapidly induced in response to various stimuli both *in vitro* and *in vivo*.

### Clec18a alleviates inflammation both in vitro and in vivo

To elucidate the functional role of Clec18a in immune regulation, we assessed inflammatory responses under conditions of Clec18a supplementation and genetic deficiency. BMDMs from Clec18a^fl/fl^LysMCre^+^ mice exhibited increased TNF-α and IL-6 production upon LPS challenge (Figure 2A and B). Myeloid-specific deficiency of Clec18a resulted in a decreased Clec18a level and elevated systemic cytokines (Figure 2C-2E) in the serum of mice six hours post-stimulation. Meanwhile, Clec18a^fl/fl^LysMCre^+^ mice also showed reduced survival rates in both the FIP and CLP sepsis models (Figure 2F and G), underscoring the critical role of myeloid-derived Clec18a in regulating inflammation. Conversely, a single dose of exogenous recombinant Clec18a (rClec18a) administration to LPS-challenged mice resulted in an elevated Clec18a in serum within six hours (Figure S2A), suppressed TNF-α and IL-6 production in a dose-dependent manner (Figure 2H), and significantly improved survival outcomes in a murine endotoxemia model (Figure 2I) and FIP model (Figure 2J). The recombinant proteins produced in HEK293 cells or *E. coli* both attenuate LPS-induced cytokine production in BMDMs (Figure S2B), suggesting glycosylation or other eukaryotic modifications of Clec18a are not essential for its function. Moreover, rClec18a also attenuated TNF-α and IL-6 levels in the serum of CpG- and poly(I:C)-challenged mice (Figure S2C), suggesting that Clec18a can broadly attenuate inflammatory signaling mediated by TLRs. This anti-inflammatory effect was recapitulated *in vitro* across murine BMDMs (Figure S2D. S2E), peritoneal macrophages (Figure S2F), and human PBMCs (Figure S2G). Furthermore, rClec18a suppressed cytokine production in BMDCs stimulated with LPS (Figure S2H), suggesting that rClec18a has immunoregulatory activity in both innate and adaptive immune cells. In macrophage, our data showed the cytokine attenuation of rClec18a lasts within 12 hours post-LPS stimulation but diminished in 24 hours (Figure S2I). In addition to exogenous supplementation, we also examined the effects of genetic overexpression on inflammation and found the overexpression of Clec18a in RAW264.7 cells further corroborated these findings. (Figure S2J and S2K).

**Figure 2.**
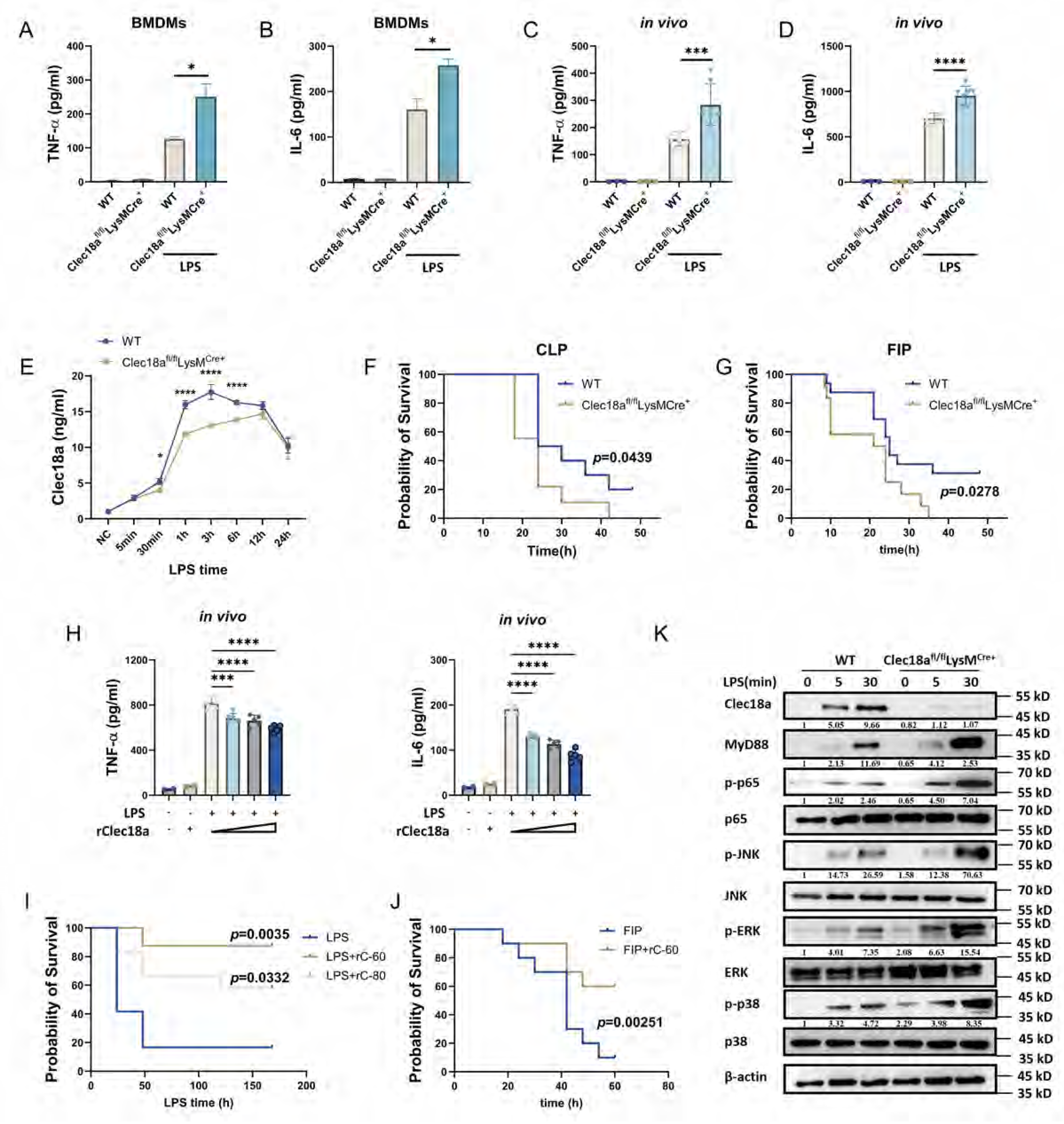
Clec18a alleviates inflammation both *in vitro* and *in vivo*. A, B: TNF-α (A) and IL-6 (B) levels in the supernatants of BMDMs from WT or Clec18a^fl/fl^LysMCre^+^ mice treated with LPS (100 ng/ml) for 6 hours (n=3). C, D: TNF-α (C) and IL-6 (D) levels in the serum of wild-type (WT) or myeloid-specific Clec18a knockout (Clec18a^fl/fl^LysMCre^+^) mice treated with *i.p.* LPS (100 ng/ml) for 6 hours (n=3-8 per group). E: Clec18a level in the serum of WT or Clec18a^fl/fl^LysMCre^+^ mice treated with LPS *i.p.* (10mg/kg) for indicated timepoints (n=4). F, G: Survival of WT or Clec18afl/flLysMCre+ mice in the cecal ligation and puncture (CLP) models (n=9 in Clec18afl/flLysMCre+ group and n=10 in littermate wild-type) (F) and fecal-induced peritonitis (FIP) (n=14 per group) (G). H: TNF-α and IL-6 levels in the serum of C57BL/6 mice treated with varying doses of mouse recombinant Clec18a (rClec18a) (20, 40, 60, 80 ng/g) and *i.p.* LPS simultaneously for 6 hours (n=3-5 per group). I: Survival of C57BL/6 mice injected *i.p.* with LPS (100 mg/kg) and rClec18a at 60 or 80 ng/g (n=16 per group). J: Survival of C57 BL/6 mice in FIP model injected *i.p.* with rClec18a at 60 ng/g (n=10 per group). K: Western blot analysis of BMDMs from WT or Clec18a^fl/fl^LysMCre^+^ mice stimulated with LPS for various durations. *p<0.05, ***p<0.001, ****p<0.0001; two-way ANOVA (A-E); Mantel‒Cox test (F,G,I,J); and one-way ANOVA (H); n=biological replicates for in vivo assay and replicate wells for in vitro assay; Data shown in all panels are representative of at least three independent experiments.

We conducted a comparative analysis of Clec18a and the classical anti-inflammatory cytokine IL-10. Temporal profiling revealed that Clec18a production precedes IL-10 production in LPS-treated mice (Figure 1F). Unlike PGE2, a previously reported inhibitory molecule,[26, 27] rClec18a does not stimulate IL-10 secretion in macrophages (Figure S2L) or in vivo (Figure S2M), suggesting that Clec18a is an early-responding negative regulator that functions independently of IL-10.

The MyD88-p65/ERK/JNK/p38 signaling pathway is activated in response to TLR4-mediated recognition. We found Clec18a attenuated TLR4-driven MyD88 activation and the phosphorylation of p65, ERK, JNK, and p38 (Figure S2N). Conversely, Clec18a-deficient BMDMs presented upregulated MyD88 expression and increased activation of these signaling nodes (Figure 2K), indicating the negative regulatory effect of Clec18a on TLR4-MyD88 signaling.

Collectively, these data establish Clec18a as a myeloid-intrinsic suppressor of inflammatory responses.

### Clec18a regulates TLR4 signaling via its receptor, Mincle

As a secretory protein, Clec18a functions through its receptor, which was unknown until now. Previous studies revealed that Clec18a can downregulate the expression of FcRγ, a critical adaptor that associates with ITAM domains in Dectin-2 family receptors (Mincle, DCAR, MCL) to initiate signaling[28] and inhibit the phagocytic activity of macrophages.[25] We speculated that molecules from the Dectin-2 family may be involved in the inhibitory function of Clec18a and performed immunoprecipitation assays to identify possible receptors of Clec18a. Surprisingly, we found that Clec18a could interact with Mincle (Figure 3A). To verify the direct interaction of Clec18a and Mincle, we performed molecular binding assays using Clec18a, Mincle and TDB, a canonical Mincle ligand. The results demonstrated that Clec18a exhibited dose-dependent Mincle binding as TDB (Figure 3B), and this interaction is independent from glycosylation, since rClec18a derived from 293T and E. Coli. showed similar affinity (Figure S3A). In addition, his-tagged BSA (same tag as rClec18a purification) and heat-inactivated rClec18a showed no interaction with Mincle, and Polymyxin B (PMB), a cationic polypeptide antibiotic that binds and abrogates the biological effects of endotoxin, co-administration didn’t influence the rClec18a-Mincle binding (Figure S3A), ruling out the endotoxin- or tag-related effects. We further applied surface plasmon resonance (SPR) assay to confirm their direct interaction. Through bidirectional assay (Mincle-Fc immobilized and rClec18a immobilized) using purified, endotoxin-free proteins, we showed that, when Mincle-Fc was immobilized with Clec18a in solution, the KD value was 548 nM (Ka of 1.3 × 10⁵ M⁻¹s⁻¹; Kd of 0.0713 s⁻¹) (Figure 3C); when Clec18a was immobilized with Mincle-Fc in solution, the KD value was 752nM (Ka of 1.18 × 10⁵ M⁻¹s⁻¹; Kd of 0.0886 s⁻¹) (Figure 3D). Thus, these results confirmed the specificity and directness of the Clec18a-Mincle interaction.

**Figure 3.**
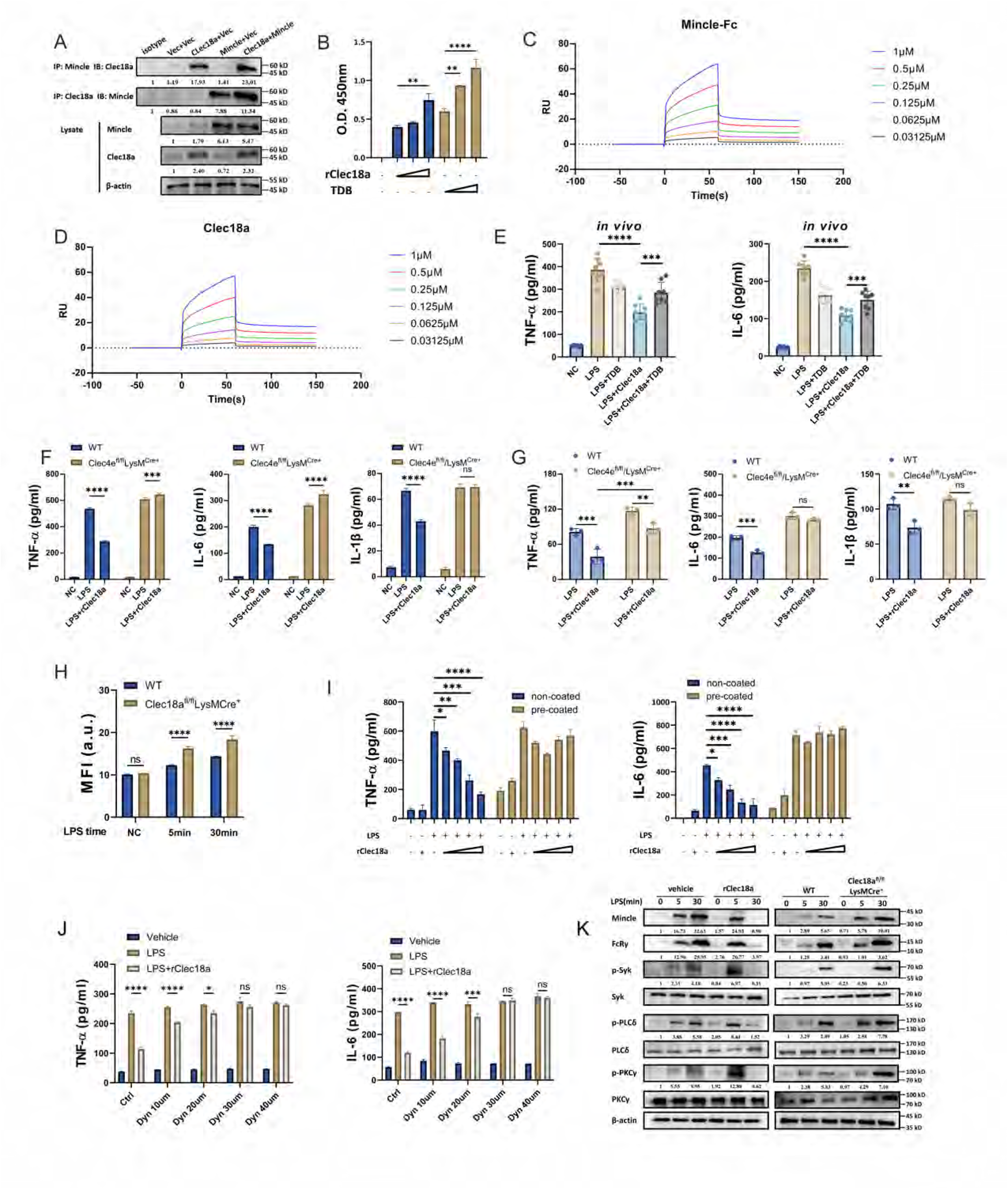
Clec18a regulates TLR4 signaling via its receptor, Mincle. A: Immunoprecipitation of Clec18a and Mincle in HEK293T cells transfected with Clec18a overexpression, Mincle overexpression, or control vectors; Lysate from Clec18a+Mincle co-transfection used in isotype lane. B: Cell-free binding assay (OD 450 nm) of rClec18a or TDB incubated with Mincle-Fc (rClec18a coated at 2.5, 5, and 10 ng/ml; TDB coated at 2.5, 5, and 10 μg/ml) (n=3). C, D: Surface plasmon resonance responses to different concentrations of Mincle-Fc (C) and rClec18a (D) during a 100-s injection. E: TNF-α and IL-6 levels in the serum of mice stimulated (injected) with LPS and rClec18a (20 ng/g) or TDB (50 μg/mouse) (pretreatment) for 6 hours (n=5-8 per group). F, G: TNF-α, IL-6 and IL-1 β levels in the supernatant of PMs (F) and in the serum (G) from Clec4e^fl/fl^LysM^Cre+^ and littermate wild-type mice and treated with rClec18a (20 ng/ml) and LPS for 6 hours (n=3). H: Flow cytometry analysis of the Mincle levels on the cell surface of BMDMs from WT and Clec18a^fl/fl^LysMCre^+^ mice treated with LPS for the indicated time points (n=4). I: TNF-α and IL-6 levels in the supernatant of PMs cultured in rClec18a-coated plates (precoated) or noncoated plates and stimulated with LPS and rClec18a for 6 hours (n=3). J: TNF-α and IL-6 levels in the supernatant of BMDMs treated with LPS+rClec18a (20 ng/ml) and Dynasore at various concentrations for 6 hours (n=3). K: Western blot analysis of BMDMs from C57BL/6 mice and WT and Clec18a^fl/fl^LysMCre^+^ mice treated with rClec18a, LPS, or a combination of both at various time points. *p<0.05, **p<0.01, ***p<0.001, ****p<0.0001, ns: not significant; one-way ANOVA (E, I) and two-way ANOVA (B, F, G, H, J); n=biological replicates for in vivo assay and replicate wells for in vitro assay; Data shown in all panels are representative of at least three independent experiments.

In investigating possible competitive binding between Clec18a and TDB, we detected mutual dose-dependent displacement between Clec18a and TDB via a molecular binding assay (Figure S3B). Coadministration of TDB also dose-dependently impaired rClec18a-mediated cytokine suppression (Figure S3C) in BMDMs and in LPS-treated mice (Figure 3E). Accordingly, TDB cotreatment reversed the rClec18a-induced suppression of MyD88, p-p65 activation and previously proposed Mincle-Syk-Card9 pathway[29] after LPS stimulation (Figure S3D). These data confirmed that Clec18a competes with TDB to bind to the receptor Mincle.

To validate the necessity of Mincle in Clec18a function, we obtained myeloid-specific Mincle knockout mice (Clec4e^fl/fl^LysM^Cre+^). Mincle deficiency abrogated the cytokine attenuation (Figure 3F) and down-regulation of MyD88-p65/ERK/JNK/p38 signaling pathway (Figure S3E) by rClec18a in LPS-stimulated BMDMs. Meanwhile, myeloid-deficiency of Mincle also reversed the cytokine attenuation of rClec18a in serum in LPS-challenged mice (Figure 3G). Blockade using anti-Mincle neutralizing antibodies showed similar reversal of Clec18a-mediated cytokine suppression in both Clec18a-overexpressing RAW264.7 cells (Figure S3F) and LPS-challenged mice (Figure S3G). Together, these results confirmed that Mincle is essential for the anti-inflammatory activity of Clec18a.

We also investigated the dynamic expression of Mincle and found that compared with LPS treatment alone, simultaneous Clec18a and LPS treatment induced rapid Mincle expression at five minutes, which could not be inhibited by Act D or CHX (Figure S3H and S3I), suggesting that the Clec18a-induced Mincle expression likely via the mobilization of pre-existing intracellular Mincle pools rather than de novo Mincle synthesis. Unlike TDB, and its analog trehalose-6,6-dimycolate (TDM), upregulates Mincle expression and promotes its translocation through mechanisms involving C/EBP *β* and MCL, respectively[30, 31, 32], Clec18a-driven Mincle membrane translocation occurred independently of MCL at 5 minutes (Figure S3J). And MCL or C/EBP *β* knockdown resulted in partial attenuation of the effects of rClec18a (Figure S3K-M), which suggests that the regulatory mechanisms through which Clec18a regulates Mincle expression and translocation are different from those of TDB/TDM.

CLRs recognize and internalize their ligands, leading to the activation of various signaling pathways.[33] Here, Clec18a deficiency elevated Mincle on cell surface (Figure 3H), while rClec18a decreased the membrane localization within 12 hours post-LPS stimulation (Figure S3N), which is similar to the regulatory window of rClec18a in cytokines (Figure S2I). Interestingly, inhibiting Mincle endocytosis through pre-coating rClec18a on plate (Figure 3I), or pre-treatment with Dynasore, a dynamin-dependent endocytosis inhibitor (Figure 3J), dose-dependently reversed the cytokine attenuation, indicating Mincle endocytosis being essential for Clec18a function. Similar to Mincle regulation, Clec18a also suppressed Mincle downstream effectors (Syk, PKCδ, PLCγ) after 30 minutes of LPS stimulation, whereas Clec18a deficiency resulted in an increase in Mincle expression and subsequent downstream signaling within 30 minutes (Figure 3K). Interestingly, the attenuation of Clec18a in Mincle signaling also relies on Mincle endocytosis, as Dynasore treatment abrogated signal attenuation at 30 minutes (Figure S3O). Thus, these data suggest that Clec18a regulates Mincle signaling by promoting Mincle endocytosis.

We subsequently excluded the possibility that other reported inhibitory intermediates associated with Mincle-Syk signaling regulation, such as SOCS1, A20,[34] SHP-1,[35] SHP-2,[36] and Cbl-b,[37] might be involved in Clec18a regulation. rClec18 led to the downregulation of Cbl-b, Cbl-c, and SHP-1 (Figure S3P). Although A20, SOCS1 and SHP-2 levels increased within five minutes, the knockdown of these molecules (Figure S3Q) did not abolish the rClec18a-mediated suppression of cytokines (Figure S3R and S3S), thus ruling out the possibility that the above molecules are involved in Clec18a-mediated immunoregulation.

Collectively, these findings establish that Clec18a regulates TLR4 signaling through Mincle.

### Clec18a induces Mincle and MyD88 degradation through Cbl-mediated ubiquitination

A previous study reported that ligands are internalized and subsequently degraded through the ubiquitin‒ proteasome system or lysosomes after binding with CLRs.[33] Our data revealed that Clec18a triggered Mincle degradation within 30 minutes. Notably, the expression and degradation of MyD88 followed a similar pattern with that of Mincle (Figure 4A), suggesting that these two molecules may be regulated by similar mechanisms. Protein degradation usually involves proteasome- or lysosome-related pathways. In our study, MG132 (an inhibitor of the proteasome), but not chloroquine (an inhibitor of lysosomes), abrogated Clec18a-mediated cytokine suppression (Figure 4B) and Mincle and MyD88 degradation in BMDMs (Figure 4C), indicating that the proteasome pathway is the primary degradation pathway. We employed the ubiquitin ligase (E3)-substrate interaction network to identify potential E3 ligases that may be involved in the ubiquitination of Mincle and MyD88. Based on the network analysis (Figure S4A and B), we knocked down candidate E3 ligases respectively to screen potential E3 ligases (data not shown) and observed that only Cbl knockdown was able to abrogate the cytokine suppression, as well as MyD88 and Mincle degradation, induced by Clec18a (Figure S4C and D). Conversely, compared with the vector control, the overexpression of Cbl in RAW264.7 cells promoted Mincle and MyD88 degradation in the presence of LPS alone and further increased their degradation and cytokine attenuation when they were cotreated with LPS and rClec18a (Figure S4E and F). In addition, co-transfection of Cbl with either Mincle or MyD88 in HEK293T cells demonstrated that Cbl promotes the degradation of both Mincle and MyD88 (Figure S4G). In myeloid-specific Cbl-knockout (Cblfl/flLysMCre+) macrophages, we observed that Cbl deficiency completely inhibited the rClec18a-mediated degradation of Mincle, MyD88, and p-p65 within 30 minutes of LPS exposure (Figure 4D), indicating that Cbl is responsible for Clec18a-mediated Mincle and MyD88 degradation. More importantly, Cbl deficiency increased Mincle expression within five minutes in the presence of LPS alone (Figure 4D), suggesting that Cbl may be involved in the endogenous Clec18a-induced degradation of Mincle. Immunoprecipitation analysis of HEK293T cells revealed that Cbl can interact with both Mincle and MyD88 (Figure 4E). Subsequent analysis of ubiquitination revealed that in LPS-stimulated Cbl-overexpressing RAW264.7 cells, rClec18a facilitated the ubiquitination of Mincle and MyD88, whereas the overexpression of Cbl further increased Clec18a-mediated Mincle and MyD88 ubiquitination (Figure S4H and I). Conversely, Cbl deficiency reversed the increase in Mincle and MyD88 ubiquitination induced by rClec18a (Figure 4F and G). Finally, through in vitro and in vivo analyses of cytokines in BMDMs and endotoxemic wild-type and Cblfl/flLysMCre+ mice, we showed that Cbl deficiency almost abrogated rClec18a-mediated cytokine suppression (Figure 4H and I) and Mincle endocytosis (Figure S4J). Thus, these findings identify Cbl as the essential E3 ligase that mediates Clec18a-induced degradation of Mincle and MyD88.

**Figure 4.**
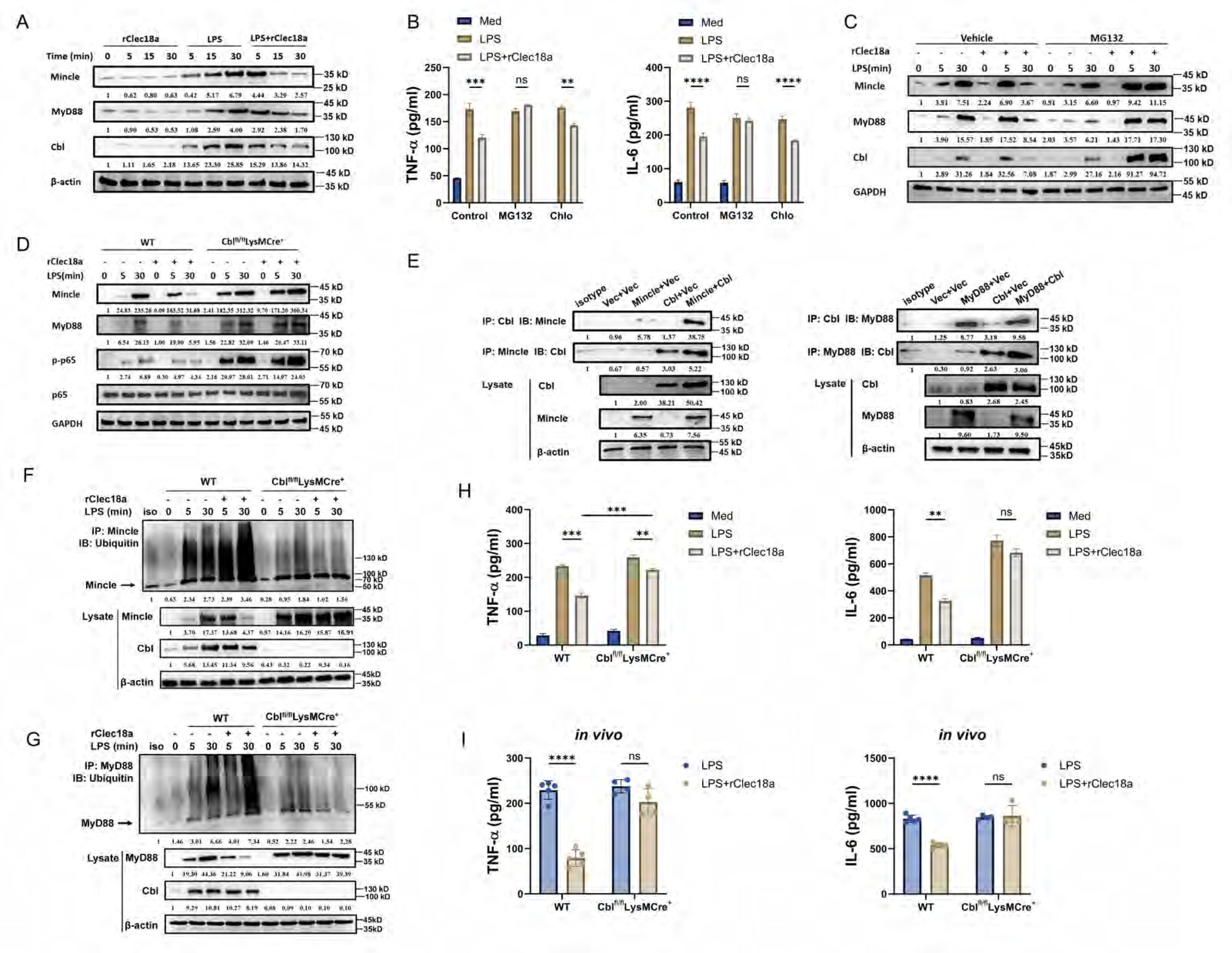
Clec18a induces Mincle and MyD88 degradation through Cbl-mediated ubiquitination. A: Western blot analysis of BMDMs from C57BL/6 mice treated with LPS, rClec18a or their combination for the indicated time points. B: TNF-α and IL-6 levels in the supernatant of BMDMs pretreated with MG-132 (20 μM) or chloroquine (20 μM) and stimulated with LPS or LPS+rClec18a for 6 hours (n=3). C: Western blot analysis of BMDMs treated with LPS and rClec18a preincubated with MG-132 or control vehicle. D: Western blot analysis of PMs from WT or Cbl^fl/fl^LysMCre^+^ mice stimulated with LPS and rClec18a for the indicated time points. E: Immunoprecipitation of Cbl with Mincle and MyD88 in HEK293T cells transfected with the indicated plasmids; Lysate from LPS-30min (WT) used in isotype lane. F and G: Western blot analysis of ubiquitin in immunoprecipitated Mincle (F) and MyD88 (G) from BMDMs of Cbl^fl/fl^LysMCre^+^ and WT mice; Lysate from LPS-30min (WT) used in isotype lane. H and I: TNF-α and IL-6 levels in the supernatant of BMDMs (n=3) (H), and in the serum of WT and Cblfl/flLysMCre+ mice (n=4 per group) (I) stimulated with LPS and rClec18a (i.p.). *p<0.05, **p<0.01, ***p<0.001, ns: not significant; two-way ANOVA; n=biological replicates for in vivo assay and replicate wells for in vitro assay; Data shown in all panels are representative of at least three independent experiments.

We also examined the potential involvement of Cbl-b, another E3 ligase from the Cbl family that has been implicated in regulating MyD88 in a previous study[37]. Knockdown of Cbl-b did not abrogate the cytokine suppression mediated by rClec18a (Figure S4K), thereby excluding the involvement of Cbl-b in the regulation of Clec18a.

Collectively, our findings suggest that Clec18a facilitates the degradation of Mincle and MyD88 via Cbl-mediated ubiquitination.

### Clec18a induces Cbl activation by regulating extracellular calcium influx

We next explored how Cbl was rapidly activated at early time points by Clec18a in the presence of LPS. Previous studies have identified an EF-hand domain in Cbl that is capable of Ca*²⁺* binding.[38] To test whether Cbl is activated via direct Ca*²⁺* binding, we first used the broad-spectrum calcium chelator BAPTA to assess Ca*²⁺* involvement in the inhibitory effect of Clec18a. BAPTA attenuated the cytokine suppression mediated by Clec18a in a dose-dependent manner (Figure 5A). Furthermore, BAPTA attenuated the ubiquitination of Mincle and MyD88 (Figure 5B and C) in five minutes induced by rClec18a. Additionally, calcium chelation also attenuated cytokine attenuation in LPS-stimulated Cbl-overexpressing RAW264.7 cells (Figure 5D). These data suggest that calcium is crucial for Cbl function. To demonstrate the direct role of calcium in Cbl activation, we utilized a cell-free system[39] to assess the ubiquitination of Cbl in the presence or absence of Ca2+ and used a well-studied calcium-binding E3 ligase, Nedd4, as a positive control. Our data indicate that the ubiquitin activity of Cbl was activated in the presence of Ca2+, with a level comparable to that of Nedd4 (Figure 5E), suggesting that calcium could activate Cbl directly. Furthermore, mutant Cbl variants with mutations in the EF-hand region (D227Q and E238S), which impairs the binding site with calcium,[38] failed to induce ubiquitination in the presence of Ca2+ (Figure 5F). Meanwhile, transfection of mutant Cbl mRNA to Cblfl/flLysMCre+ macrophage abolished the Mincle and MyD88 ubiquitination induced by rClec18a (Figure 5G). These results indicate that Ca2+ interacts with the EF-hand region to induce Cbl activation. Intriguingly, calcium imaging analysis revealed that, although the signal was decreased compared with that of LPS alone, exposure to rClec18a and LPS resulted in an earlier increase in the intracellular calcium signal, whereas exposure to rClec18a alone resulted in no calcium flux (Figure 5H).

**Figure 5.**
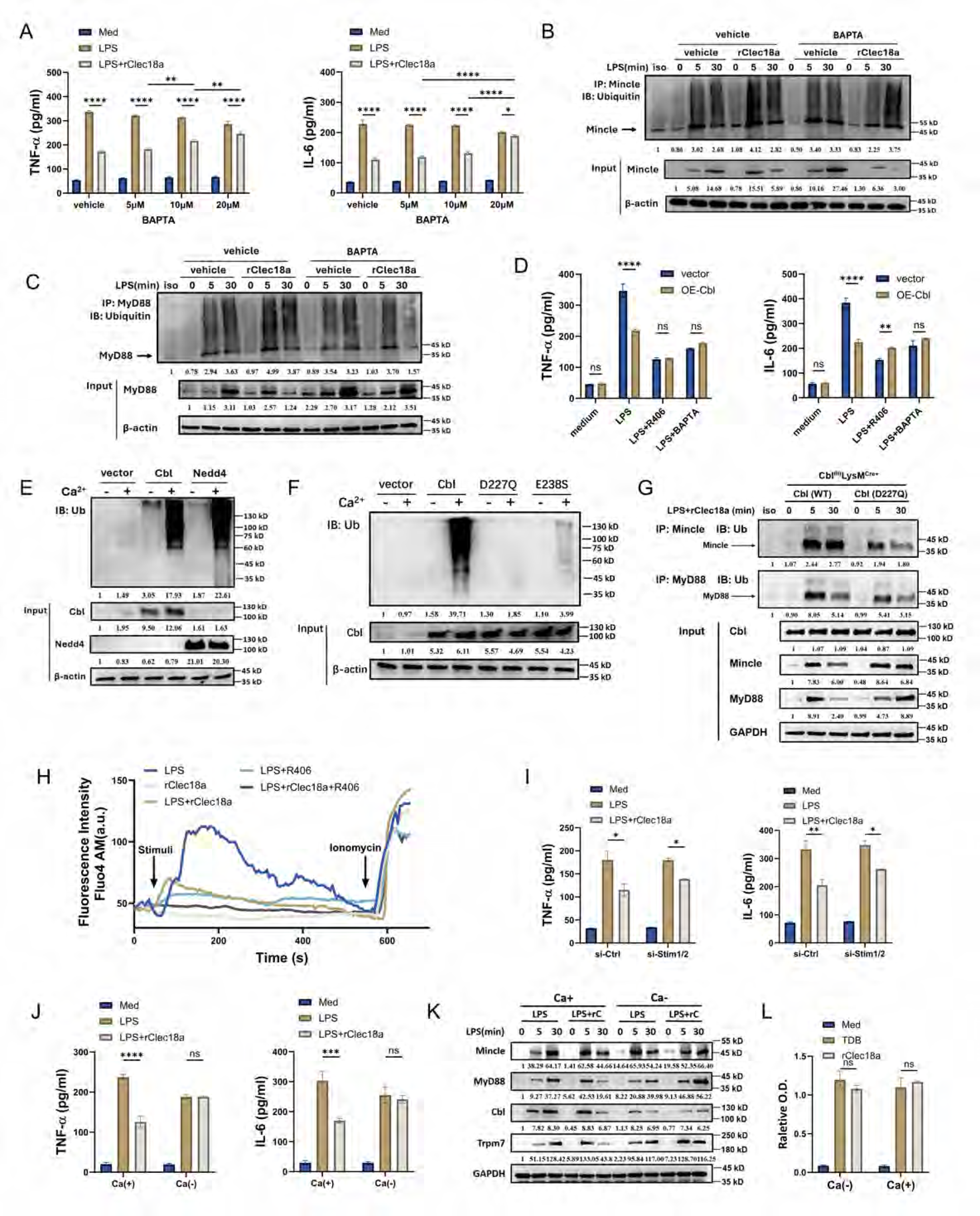
Clec18a induces Cbl activation by regulating extracellular calcium influx. A: TNF-α and IL-6 levels in the supernatant of BMDMs pretreated with BAPTA (calcium chelator) (5, 10, 20 μM) at indicated concentrations for 30 minutes and then stimulated with LPS and rClec18a for 6 hours (n=3). B, C: Western blot analysis of ubiquitin of immunoprecipitated Mincle (B) and MyD88 (C) in PMs pretreated with BAPTA (20 μM) and then stimulated with LPS or LPS+rClec18a; Lysate from LPS-30min (vehicle) used in isotype lane. D: TNF-α and IL-6 levels in the supernatant (n=3) of RAW264.7 cells overexpressing Cbl and treated with the indicated reagents (BAPTA, 20 μM or R406, 2 μM). E: Ubiquitin activity assay of immunoprecipitated Cbl, Nedd4, and vector plasmid from HEK293 cells in a cell-free mixture with (Ca^2+^+) or without CaCl_2_ (Ca^2+^-), input represents Western blot analysis of Cbl and Nedd4 from immunoprecipitation, Nedd4 as positive control. F: Ubiquitin activity assay of immunoprecipitated Cbl, EF-hand-mutant Cbl (D227Q or E238S), and vector plasmid from HEK293 cells in a cell-free mixture with (Ca^2+^+) or without CaCl_2_ (Ca^2+^-), input represents Western blot analysis of Cbl from immunoprecipitation. G: Western blot analysis of ubiquitin of immunoprecipitated Mincle and MyD88 in PMs from Cbl^fl/fl^LysMCre^+^ mice transfected with wild-type or mutant (D227Q) Cbl mRNA stimulated with LPS or LPS+rClec18a; Lysate from 30min (Cbl WT) used in isotype lane. H: Calcium flux imaging by confocal microscopy in PMs treated with LPS, LPS+rClec18a, or rClec18a with or without R406. I: TNF-α and IL-6 levels in the supernatant of PMs with Stim1/2 knocked down for 48 hours and stimulated with LPS for 6 hours (n=3). J: TNF-α and IL-6 levels in the supernatant (n=3) of BMDMs stimulated with LPS or LPS+rClec18a (six hours for cytokine assay) in Ca^2+^-containing or Ca^2+^-free medium. K: Western blot analysis of BMDMs cultured in medium with (Ca^2+^+) or without CaCl_2_ (Ca^2+^-) and treated with LPS or LPS+rClec18a for the indicated durations. L: Cell-free binding assay (O.D. 450nm) of rClec18a or TDB incubated with Mincle-Fc in Ca2+-containing or Ca2+-free medium (n=3). **p<0.01, ***p<0.001, ****p<0.0001, ns: not significant; two-way ANOVA; n=biological replicates for in vivo assay and replicate wells for in vitro assay; Data shown in all panels are representative of at least three independent experiments.

LPS-induced calcium mobilization originates from both extracellular influx through transient receptor potential channels (TRPCs), such as Trpm7,[40, 41] and from the endoplasmic reticulum through a store-operated Ca2+ entry (SOCE) mechanism.[42] We wondered which source of calcium was essential for Clec18a function. Unexpectedly, the knockdown of Stim1/2, crucial modulators of SOCE, had a limited influence on the cytokine suppression mediated by rClec18a (Figure 5I). However, in calcium-deficient culture medium, Clec18a-mediated cytokine suppression and the degradation of Cbl, MyD88 and Mincle were almost completely abrogated (Figure 5J and K). Although a previous study indicated that calcium may inhibit the ligand recognition of Mincle,[43] our data revealed that calcium deficiency had no effect on the Clec18a-Mincle interaction (Figure 5L). Thus, these results indicate that Clec18a induces Cbl activation through extracellular calcium influx.

### Trpm7-derived calcium is responsible for Clec18a-mediated Cbl activation

Previous studies revealed that the early increase in cytosolic Ca2+ induced by LPS could result from influx through membrane ion channels, such as transient receptor potential melastatin-like 7 (Trpm7), which is activated in a CD14 endocytosis-dependent manner.[40, 42] The reliance on endocytosis and early calcium flux prompted us to investigate the potential involvement of Trpm7 in Clec18a-mediated calcium mobilization. We found that the knockdown of Trpm7 negated the cytokine suppression (Figure 6A), degradation of Mincle, MyD88, and Cbl (Figure 6B), and ubiquitination of Cbl (Figure 6C) induced by Clec18a, indicating that Trpm7 is crucial for Cbl activation. To determine whether Clec18a regulates the functional activity of Trpm7 channels on the plasma membrane, we performed whole-cell patch-clamp electrophysiology to record Trpm7-like currents in macrophages followed previously established protocols[44, 45]. As shown in the current-voltage (I-V) relationship, wild-type macrophages elicited robust outwardly rectifying currents characteristic of Trpm7 (Figure 6D). In contrast, these currents were severely abrogated in Clec18afl/flLysMCre+ macrophages, resulting in a significant reduction in peak current amplitude compared to wild-type controls (Figure 6D). These results suggested Clec18a is essential for channel activity of Trpm7.

**Figure 6.**
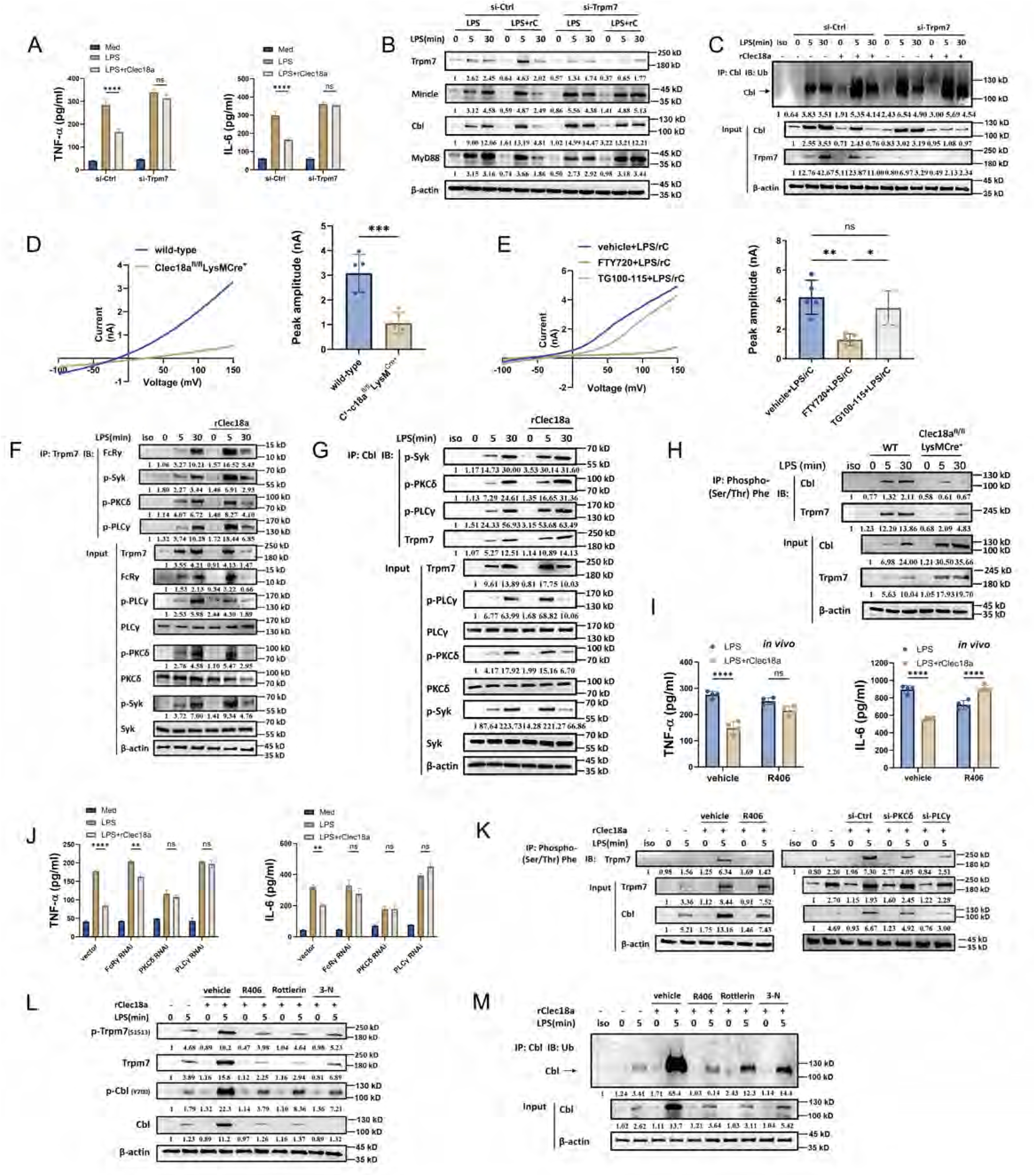
Trpm7-derived calcium is responsible for Clec18a-mediated Cbl activation. A: TNF-α and IL-6 levels in the supernatant (n=3) of PMs with Trpm7 knockdown and then stimulated with LPS or LPS+rClec18a for 6 hours. B: Western blot analysis of PMs with Trpm7 knockdown and then stimulated with LPS or LPS+rClec18a for indicated timepoints. C: Western blot analysis of ubiquitin of immunoprecipitated Cbl in PMs with Trpm7 knockdown and treated with LPS or LPS+rClec18a at the indicated time points. D: Representative whole-cell patch-clamp recordings of Trpm7 currents in LPS-stimulated wild-type and Clec18a^fl/fl^LysM^Cre+^ macrophages. The left panel displays representative current-voltage (I-V) traces showing the characteristic outwardly rectifying currents. The right panel shows the quantification of peak current amplitudes measured at +150 mV (n=5 cells per group); Data are presented as mean ± SEM. E: Clec18a^fl/fl^LysM^Cre+^ macrophages were treated with vehicle, the Trpm7 channel blocker FTY720, or the Trpm7 kinase inhibitor TG100-115 prior to LPS+rClec18a stimulation. The left panel displays representative current-voltage (I-V) traces showing the characteristic outwardly rectifying currents. The right panel shows the quantification of peak current amplitudes measured at +150 mV (n=5 cells per group); Data are presented as mean ± SEM. F, G: Immunoprecipitation of Trpm7 (F) and Cbl (G) with FcRγ, p-Syk, p-PKCδ and p-PLCγ in PMs treated with LPS or LPS+rClec18a at the indicated time points. Lysate from LPS-30min (vehicle) used in isotype lane. H: Immunoprecipitation of phosphor-(Ser/Thr) Phe from Cbl and Trpm7 in PMs from WT and Clec18a^fl/fl^LysMCre^+^ mice treated with LPS at the indicated time points. Lysate from LPS-30min (WT) used in isotype lane. I: TNF-α and IL-6 levels in the serum of mice treated with the Syk inhibitor R406 (5 mg/kg) or the control vehicle and stimulated with LPS or LPS+rClec18a for 6 hours (n=4 per group). J: TNF-α and IL-6 levels in the supernatant of PMs with FcRγ, PKCδ or PLCγ knockdown and stimulated with LPS or LPS+rClec18a for 6 hours (n=3). K: Immunoprecipitation of phosphor-(Ser/Thr) Phe from Trpm7 in PMs pretreated with R406 or with PKCδ and PLCγ knockdown for 48 h and stimulated with LPS and rClec18a for the indicated time points. Lysate from LPS-5min (vehicle) used in isotype lane. L: Western blot analysis of BMDMs pretreated with the indicated inhibitors and stimulated with LPS and rClec18a for the indicated durations. M: Ubiquitination assay of immunoprecipitated Cbl in PMs pretreated with inhibitors of Syk (R406, 2 μM), PKCδ (Rottlerin, 20 μM) or PLCγ (3-N, 30 μM) and stimulated with LPS or LPS+rClec18a for the indicated durations. Lysate from LPS-5min (vehicle) used in isotype lane. *p<0.05, **p<0.01, ***p<0.001, ****p<0.0001, ns: not significant; Student’s *t* test (D), one-way ANOVA(E), two-way ANOVA (I, J); n=biological triplicates; n=biological replicates for in vivo assay and replicate wells for in vitro assay; Data shown in all panels are representative of at least three independent experiments.

Trpm7 functions through both its ion channel and kinase activities.[46, 47] We found that the pharmaceutical inhibition of the channel activity of Trpm7 by FTY720[40, 41] resulted in an attenuation of cytokines similar to that of BAPTA (Figure S5A), whereas inhibitors of kinase activity by TG100-150[48, 49] had no effect (Figure S5B). Moreover, the ubiquitination of Cbl was attenuated by FTY720 (Figure S5C) in five minutes post-LPS/rClec18a administration, suggesting that the ion channel activity of Trpm7 is crucial for Cbl activation.

Indeed, through patch-clamp assay, treatment with FTY720 recapitulated the phenotype observed in Clec18a-deficient cells, effectively suppressing the outwardly rectifying currents compared to the vehicle control (Figure 6E). Conversely, the Trpm7 kinase inhibitor TG100-115 did not significantly alter the channel current amplitude (Figure 6E), suggesting the observed effect is specific to the channel pore function.

Taken together, these results indicate that Clec18a activates Cbl through Trpm7-derived extracellular calcium influx.

We next explored how Clec18a regulates Trpm7 expression. We found that the expression of Trpm7 exhibited a pattern similar to that of Mincle, Cbl, and MyD88 in the presence of Clec18a (Figure S5D). Both the proteasome inhibitor MG-132 and the endocytosis inhibitor dynasore effectively inhibited the downregulation of Trpm7 induced by rClec18a within 30 minutes following LPS stimulation (Figure S5E and F). Meanwhile, Trpm7 interacted with MyD88, along with Mincle and Cbl (Figure S5G). Cbl deficiency reversed the Clec18a-induced down-regulation of Trpm7, along with Mincle and MyD88 (Figure S5H). The ubiquitin ligase (E3)-substrate interaction network predicts Trpm7 as a substrate of Cbl (Figure S5I). And Clec18a-mediated Trpm7 down-regulation was dependent on calcium (Figure 5K). These results suggest that Trpm7 may undergo degradation via ubiquitination regulated by Clec18a-induced Cbl activation, and, thus, results in the termination of early calcium flux.

Previous research has demonstrated that the cellular Ca2+ flux mediated by Trpm7 can be activated by PLCγ activity,[50] and Cbl phosphorylation is crucial for its ubiquitination activity.[51, 52, 53] To gain deeper insight into Clec18a-mediated Trpm7 and Cbl activation, we investigated the role of the Syk-PLCγ signaling, a downstream pathway of Mincle. Clec18a activated Syk-PKCδ-PLCγ signaling within 5 min (Figure 3K), induced the interaction of these molecules with Trpm7 (Figure 6F) and Cbl (Figure 6G), and promoted the phosphorylation of both molecules (Figure S5J). Meanwhile, Clec18a deficiency decreased their phosphorylation (Figure 6H) and largely suppressed interactions between Trpm7 and p-Syk/p-PLCγ (Figure S5K). Functionally, inhibiting Syk using R406 completely abolished the early calcium influx (Figure 5H) induced by Clec18a, which, in turn, reversed the cytokine suppression of rClec18a at six hours post-LPS stimulation both in vitro (Figure S5L) and in vivo (Figure 6I). Knockdown of FcRγ, PKCδ or PLCγ (Figure S5M-O), or pharmaceutical inhibition of these molecules, partially reversed Clec18a-mediated cytokine suppression (Figure 6J) and Trpm7/Cbl phosphorylation (Figure 6K, L) [54, 55], consistent with a contributory role for the Syk-PKCδ-PLCγ axis, though the concomitant reduction in basal LPS responses upon PKCδ knockdown precludes definitive conclusions regarding its specific requirement in Clec18a signaling. Moreover, pharmaceutical inhibition of these molecules also attenuated the ubiquitination of Cbl (Figure 6M). Thus, our results suggested that Clec18a may activate Trpm7 and Cbl through Syk-PLCγ signaling.

Taken together, these findings indicate that Clec18a activates Cbl through Trpm7-derived calcium.

### Clec18a promotes MyD88-Mincle-Cbl complex formation

The synchronous expression and simultaneous interaction with Cbl suggest the possibility of complex formation involving MyD88, Mincle, Trpm7 and Cbl. Indeed, we found that MyD88, Mincle and Cbl intrinsically interacted upon LPS ligation and that rClec18a further promoted their interaction (Figure 7A), whereas in Clec18a-deficient macrophages, the interaction between Cbl, MyD88 and Mincle, although largely compromised, was still observed (Figure 7A), suggesting that LPS inherently promoted their interactions, whereas Clec18a critically enhanced the mutual binding of the complex that established the molecular basis for Clec18a functions. A GST/His pulldown assay confirmed that MyD88, Mincle and Cbl could both directly interact (Figure 7B), indicating their intrinsic interaction. However, the interaction of Trpm7 with the Mincle-MyD88-Cbl complex was crucially dependent on Clec18a, as Clec18a deficiency reversed the interaction of Trpm7 with these three molecules (Figure 7A).

**Figure 7.**
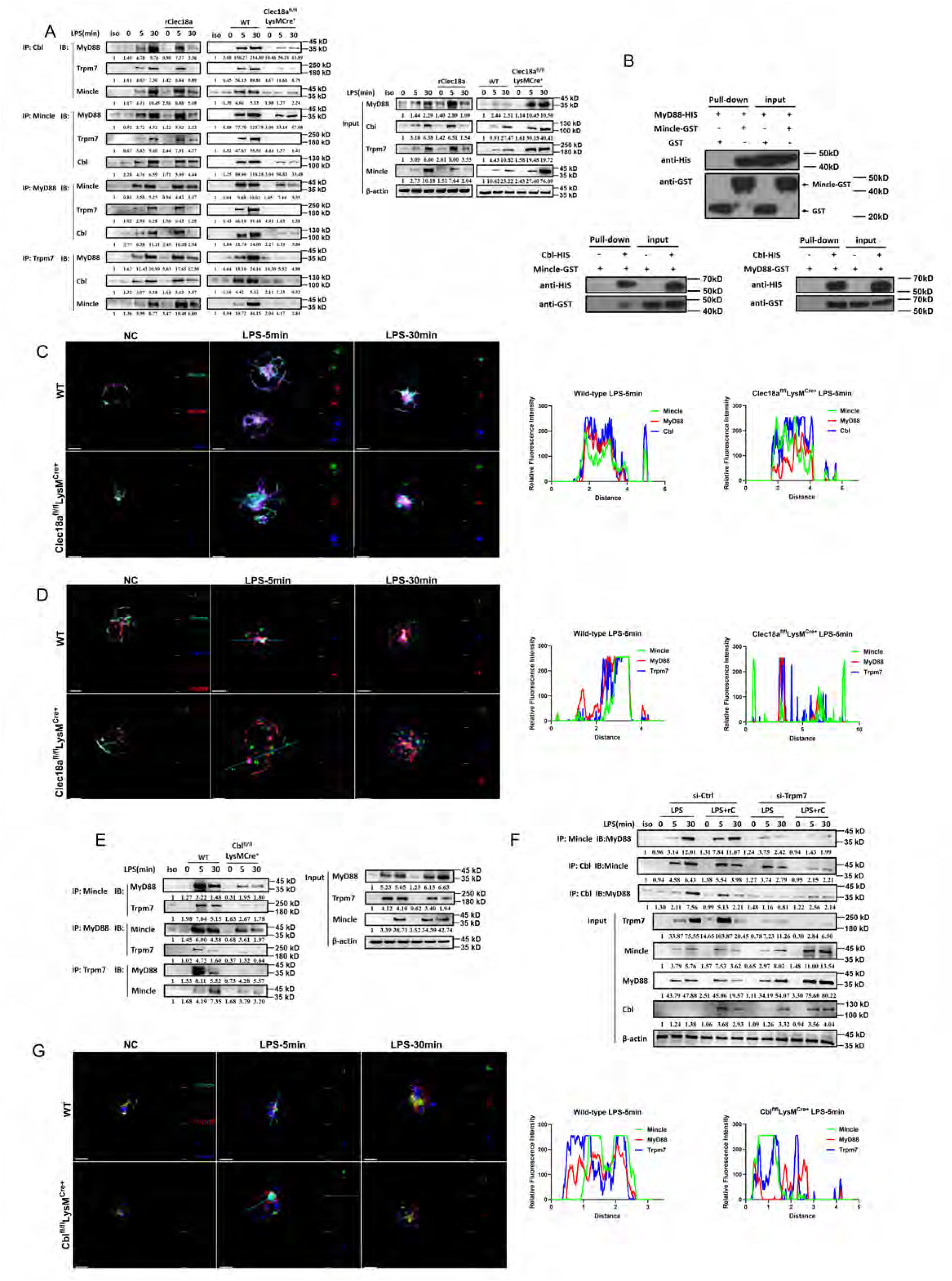
Clec18a promotes MyD88-Mincle-Cbl complex formation. A: Immunoprecipitation of MyD88, Mincle, and Cbl in PMs from C57BL/6 mice or WT and Clec18a^fl/fl^LysMCre^+^ mice stimulated with LPS or LPS+rClec18a at the indicated time points. Lysate from LPS-30min (WT) used in isotype lane. B: Pulldown assay of MyD88, Mincle, and Cbl purified using a glutathione S-transferase (GST) or histidine (HIS) tag. C, D: Confocal microscopy images and colocalization quantification (dotted line) of BMDMs from Clec18a^fl/fl^LysMCre^+^ and WT mice stimulated with LPS at the indicated time points (Bar=5μm). E: Immunoprecipitation of MyD88 and Mincle in PMs from WT and Cbl^fl/fl^LysMCre^+^ mice stimulated with LPS or LPS+rClec18a at the indicated time points. Lysate from LPS-30min (WT) used in isotype lane. F: Immunoprecipitation of MyD88, Mincle, and Cbl in PMs with Trpm7 knockdown and stimulated with LPS or LPS+rClec18a at the indicated time points. Lysate from LPS-30min (si-Ctrl) used in isotype lane. G: Confocal microscopy images and colocalization quantification (dotted line) of BMDMs from Cbl^fl/fl^LysMCre^+^ and WT mice stimulated with LPS at the indicated time points (Bar=5μm); Data shown in all panels are representative of at least three independent experiments.

Clec18a-mediated complex formation was also confirmed by confocal microscopy (Figure 7C). During the early phase of LPS stimulation (five minutes), Clec18a promoted the colocalization of Cbl, MyD88, and Mincle in wild-type macrophages. In contrast, this interaction was diminished in Clec18a-deficient macrophages (Figure 7C). Similarly, thirty minutes after LPS stimulation, the colocalization of these proteins was further increased, whereas in Clec18a-deficient macrophages, their interaction was suppressed (Figure 7C). The colocalization of Trpm7 with Mincle and MyD88 was consistent with the immunoprecipitation results. In wild-type macrophages, Trpm7 quickly colocalized with Mincle and MyD88. However, at both five and thirty minutes after LPS stimulation, Trpm7 remained distributed throughout the cells of Clec18a-deficient macrophages (Figure 7D). Taken together, these results indicated that LPS stimulated the Mincle-MyD88-Cbl interaction and that Clec18a further enhanced these interactions through Trpm7.

We then asked about the possible central role of Cbl in such dependency. In Cbl-deficient macrophages, the Clec18a-promoted Mincle-MyD88 interaction at five minutes was attenuated (Figure 7E). Moreover, the interaction of Trpm7 with these two molecules was also decreased (Figure 7E). These results indicated that the enhanced MyD88-Mincle interaction, as well as the interaction of Trpm7 with the complex mediated by Clec18a, was dependent on Cbl. This observation was further confirmed by the knockdown of Trpm7, which suppressed Cbl activation, attenuating the formation of the Cbl-Mincle-MyD88 complex (Figure 7F). These data indicated that Clec18a promoted the formation of Mincle-MyD88-Cbl-Trpm7 complex by activating Cbl. In Cbl-deficient macrophages, Mincle, MyD88 and Trpm7 were separated five minutes after LPS stimulation. Moreover, Trpm7 colocalized with Mincle and MyD88 in Cbl-deficient macrophages at both five and thirty minutes after LPS challenge (Figure 7G), which is in accordance with the immunoprecipitation results (Figure 7E), suggesting that Cbl was responsible for the early formation of the complex and especially for Trpm7 mobilization.

Collectively, these results indicate that Clec18a promotes early formation and activation of the MyD88-Mincle-Cbl complex through Trpm7, which facilitates subsequent ubiquitination-mediated degradation.

## DISCUSSION

Our study showed an elevated expression of Clec18a associated with multiple inflammatory settings, including sepsis, bacterial and viral infection, and autoimmune diseases, suggesting potential broad-spectrum regulatory function in inflammation. Clec18a responds quickly to multiple stimuli and suppresses the immune response in the early phase, adding new evidence of the regulatory mechanism in the early phase of immune responses. Unlike induced negative regulatory cytokines such as IL-10, basally stored Clec18a enables rapid integration and suppression of inflammatory signaling molecules within minutes of stimulus onset. This immediate action imposes a critical "checkpoint" on the initiation of innate immune responses, serving as a frontline mechanism to constrain excessive inflammation at its earliest stage and avert pathological hyperactivation that may otherwise lead to tissue damage or systemic disorders. Meanwhile, Clec18a functions likely in the early phase of inflammation, while it shows limited impact on down-stream molecules 12 hours after the LPS stimulation (Figure S5P). Therefore, the present study reveals a previously unrecognized intrinsic regulatory role of Clec18a in immune homeostasis.

We identified Mincle as the functional receptor for Clec18a. Compared with Mincle’s canonical ligand TDB, Clec18a exhibits a higher binding affinity for Mincle, thereby providing an additional mechanistic explanation for its ability to sequester pro-inflammatory Mincle ligands. It should also be noted that the active, intracellular mechanism discovered in the present study that is different from the conventional proinflammatory effects of Mincle activation[56, 57, 58]. TLR activation triggers Clec18a production and subsequent binding with Mincle, which promotes the formation of the MyD88-Mincle-Cbl complex and subsequent Mincle and MyD88 degradation mediated by Cbl, which bridged an intrinsic TLR‒TLR crosstalk. This negative immune regulatory mechanism of Clec18a is distinct from canonical inhibitory pathways involving SHP-1[28, 35] or other inhibitory molecules[16, 34, 35, 36, 37, 59] and from RAF1-dependent DC-SIGN or SHP-1/2 activation, which is usually triggered by dual activation of two signaling pathways.[17, 18, 19, 20, 21, 60, 61] The present study not only identified a new ligand for Mincle but also demonstrated that different ligands trigger various signaling pathways, indicating the dual role of Mincle in pathogen recognition and immune regulation. However, future studies are still needed to identify the recognition or binding region of Clec18a to Mincle with physiologically relevant affinity.

Clec18a promoted the interaction of Cbl with Mincle and MyD88 and induced complex formation and degradation, while Cbl deficiency inhibited this process. Furthermore, our data revealed a unique mechanism of Cbl activation that requires both phosphorylation and direct calcium binding. Previous studies have well-characterized the Cbl phosphorylation by Syk and downstream signaling that are crucial for ubiquitination activity [52, 54, 62], which is in accordance with our observation that Syk inhibition reversed effects of Clec18a both in cytokines and downstream signaling. Yet, our study also uncovered the calcium binding to the EF-hand motif being require for ubiquitination induction by Cbl. Although Cbl overexpression (Figure S5Q) or rClec18a treatment (Figure S5J) induced Cbl phosphorylation, it was only activated in the presence of Ca2+ (Figure 4D). Inhibition of Trpm7-originated Ca2+ abolished Cbl ubiquitination. Moreover, mutation of D227Q or E238S in the calcium-binding EF-hand motif impaired the ability of Cbl to induce ubiquitination (Figure 5F, G). This finding explained a previously reported observation that Cbl binding to TRAF6 could be antagonized by TRPM7-induced Ca2+ signaling,[63] as this may be a result of Ca2+-mediated Cbl activation and the subsequent interaction between Cbl and the MyD88-Mincle complex. However, it should be noted that Trpm7 inhibitors and BAPTA (BAPTA-AM) may have off-target effects and can influence many Ca*²⁺*-dependent processes. These inhibitor-based results are considered supportive rather than definitive. Given that global Trpm7 knockout results in embryonic lethality[64] and impaired cell proliferation[65], future studies using inducible conditional knockout models are warranted to provide definitive genetic evidence. However, our study expands the knowledge of the function and activating mechanism of Cbl and provides a basis for the development of pharmaceuticals that regulate Cbl function to treat both inflammatory and autoimmune diseases.

Unlike the previously reported proinflammatory property of Trpm7,[40, 63] Clec18a-activated Trpm7 exhibits anti-inflammatory properties through Cbl-mediated MyD88-Mincle degradation. In addition, consistent with previous findings that the Syk-PKC kinase regulates the activity of Trpm channels,[50, 66, 67, 68] our data revealed that Clec18a promoted the interaction between Syk/PKC *δ* /PLC *γ* and Trpm7, resulting in increased phosphorylation. When we analyzed Trpm7 expression, we found that the expression pattern positively correlated with that of Cbl and Mincle. Moreover, Clec18a promoted the interaction of Trpm7 with Cbl, whereas Cbl deficiency abrogated Clec18a-mediated Trpm7 degradation. Therefore, these data indicate that Trpm7 may be ubiquitously degraded through Cbl. Our data also provide new insights into the mechanisms through which CD14 regulates Trpm7 function, which was not fully understood before [40]. A possible explanation is that inactivation of CD14 may inhibit Clec18a production through CD14/MD2-mediated JNK/AP-1 activation,[69] thus impairing Trpm7 function. Our study revealed diverse properties and regulatory roles of Trpm7 in early innate immune responses.

We uncovered Clec18a-mediated formation of the MyD88-Mincle-Cbl complex, establishing a crosstalk of TLR-CLR in the early stages of the immune responses and a structure basis for the anti-inflammatory effects. Cbl, which is activated by Clec18a, plays a central role in complex formation. Previous studies have reported Cbl functions through E3 ligase activity, yet our data suggest that Cbl may function as a scaffold protein that couples TLR and CLR signaling. Although MyD88, Mincle and Cbl slightly interacted with one another without Clec18a stimulation, its ubiquitination and degradation were inhibited by Cbl deficiency. Interestingly, we discovered that Trpm7 plays an indispensable role in the formation of the MyD88-Mincle-Cbl complex induced by Clec18a, indicating that Trpm7 functions as a “molecular switch” in Clec18a regulation. Clec18a deficiency completely inhibited the interaction of Trpm7 with the other three molecules. Therefore, Clec18a-induced activation of Cbl and complex formation might be enabled by the spatial proximity of Trpm7 to Cbl. This observation is similar to the finding that Trpm7-dependent calcium influx promoted TLR4 endocytosis.[40] Thus, our data demonstrate that Clec18a-induced Trpm7 activation triggers the formation and degradation of the MyD88-Mincle-Cbl complex, bridging TLR and CLR signaling.

There are several limitations in the present study. Firstly, since potential incomplete specificity and off-target effects of the pharmacological tools, as well as knockdown efficiency, interpretation based on these results should be cautious and require further validation. Secondly, the human cohort in the present study is relatively small, limiting statistical power and generalizability. And our mechanistic conclusions rely primarily on murine experiments, while human data remains preliminary. Future studies should validate these observations in larger cohorts and in ex vivo primary human macrophages to test the conservation of the proposed mechanism.

In summary, the present study highlights an intrinsic secretory anti-inflammatory soluble C-type lectin, Clec18a, which is expressed in multiple immune cells in response to diverse harmful stimuli. Our results revealed its specific receptor, Mincle, and the subsequent recruitment and activation of the E3 ligase Cbl via the Trpm7-Ca^2+^ axis. This finding provides new insight into the negative regulation of the innate immune response, expanding the knowledge of CLR-TLR intrinsic regulation. In light of its expression in multiple cells in response to diverse stimuli, Clec18a could represent a potential therapeutic target pending further validation in human samples and demonstration of safety/efficacy.

## METHODS

### Animal model

C57BL/6J, Clec18afl/flLysMCre+, Clec4efl/flLysMCre+ and Cblfl/flLysMCre+ mice were obtained from Modelorg, Inc. (Shanghai, China). Myeloid-specific Clec18a and Cbl knockout mice were generated by crossing Clec18afl/fl or Cblfl/fl mice with LysM-Cre mice on a C57BL/6J background. The animals were housed under controlled conditions at a temperature of 18–22°C with a relative humidity of 50–60% and a 12-hour light‒dark cycle. The animals had free access to standard food and water. Ethical approval for the animal experiments was obtained from the Ethical Committee of Changhai Hospital following the Experimental Animal Regulations of Naval Medical University. To induce peritonitis in mice, lipopolysaccharide (LPS) (Escherichia coli O111:B4, Sigma, China) (10 or 40 mg/kg) diluted in 500 µl of phosphate-buffered saline (PBS) was intraperitoneally injected. To establish the cecum ligation and puncture (CLP) model, anesthesia was induced using 1.5-2% sevoflurane mixed with oxygen and an anesthetic evaporator (Drager, Germany). A midline incision measuring 10 mm was made to expose the cecum, which was ligated approximately 1 cm before being punctured at its distal end using a 22-G needle. The fecal-induced peritonitis (FIP) model was established as previously described.[70] Briefly, feces were collected from the cecal contents of C57BL/6J mice and filtered through a 100 μm cell strainer before centrifugation at 3000×g for 25 minutes at 4°C. The sediment was resuspended in PBS to a final concentration of approximately 100 mg/ml, which was then intraperitoneally injected at a dose equivalent to 0.75 mg/g body weight. All experimental mice were co-housed from weaning through the entirety of infection experiments to ensure identical environmental and microbiome exposures across genotypes.

### Gene expression omnibus (GEO) datasets analysis

Gene expression matrices from GEO datasets (GSE14905, GSE15573, GSE20346, GSE38941, GSE137340, GSE26440, GSE57065 and GSE65682) were preprocessed using standardized bioinformatic pipelines. Unannotated probes and genes with >50% missing values (expression = 0 treated as NA) were filtered. Multi-probe mappings were resolved by averaging expression values per gene symbol, followed by zero-imputation of residual NAs. Normalized expressions of merged sepsis datasets of GSE137340, GSE26440, GSE57065 and GSE65682 were presented in Table S1, a total of 1070 sample were included in the analysis. Datasets exhibiting non-normal distributions (assessed via kernel density plots) underwent gene-wise Z-score normalization to preserve relative expression patterns across samples. Differential expression of Clec18a between pre-defined case/control cohorts was evaluated using independent samples t-tests on normalized expression values. Kinetic analysis of Clec18a from GEO57065 dataset was performed with GEO2R’s default limma pipeline (R package v3.48.3). The GSEA functional enrichment analysis was conducted using the clusterProfiler package in R. The functions with a p-value less than 0.05 were selected as significant functions and visualized.

### PBMC isolation and serum collection

In analyzing Clec18a expression from PBMCs and serum, we utilized samples derived from cohort that approved by the Ethics Committee of Changhai Hospital, Naval Medical University (Approval No. CHEC2018-164) and registration with the China Clinical Trials Registry (ChiCTR1800015714). Written informed consent was obtained from all participants. Patients with infection were defined as those with a positive blood culture result at the time of sample collection; the baseline characteristics of the patients are presented in Table S2. PBMCs were isolated using standard density gradient centrifugation with Ficoll (Sigma, USA) and cultured in RPMI-1640 (HyClone, USA) for further experiments. CD14^+^ monocytes were isolated by using human CD14 MicroBeads (Miltenyi Biotec, China) according to the manufacturer’s instructions. Serum was isolated by centrifugation at 3000 × g for 15 minutes and analyzed for further experiments.

### Recombinant Clec18a protein

In this study, recombinant human and mouse Clec18a was purchased as a commercial preparation from CUSABIO (CSB-EP005521HU; CSB-BP759740MO; E. coli or HEK293–expressed; N-terminal 6×His-SUMO tag;). Importantly, according to the manufacturer’s quality control documentation, each recombinant protein undergoes a standardized QC process that includes endotoxin removal and endotoxinection by the LAL method, with an acceptance criterion of <0.05 EU/μg. The manufacturer further indicates that endotoxin levels are reported in the lot-specific certificate of analysis (COA). Consistent with these specifications, the lot used in our experiments was accompanied by a COA indicating endotoxin levels within the manufacturer’s criteria.

### Macrophage or dendritic cell

To isolate peritoneal macrophages (PMs), the mice were intraperitoneally injected with thiocyanate (BD, USA), and peritoneal lavage fluid was collected three days after injection. The PMs were resuspended at a concentration of 2∼4×10^6^ cells/ml and cultured in RPMI 1640 medium supplemented with 10% fetal bovine serum (FBS) (Gibco, USA). To isolate bone marrow-derived macrophages (BMDMs), bone marrow was collected from the femoral tissues of mice flushed with 3 ml of normal saline (NS). After red blood cell lysis, the bone marrow cells were resuspended at a concentration of 2–4×10^6^ cells/ml in Dulbecco’s modified Eagle’s medium (DMEM) supplemented with 10% FBS and 30 ng/ml GM-CSF. The culture medium was changed every two days, and after 5–6 days of culture, the cells were used for further experiments. LPS was added to the cells at a concentration of 100 ng/ml to stimulate the macrophages for the indicated times. Bone marrow-derived dendritic cells (BMDCs) were prepared following the same protocol as that used for BMDM isolation and resuspended in RPMI-1640 culture medium supplemented with 10% FBS, 20 ng/ml GM-CSF and 10 ng/ml IL-4. After six days of culture, the suspended cells were collected and subjected to further experiments. CD11c^+^ DCs were isolated with a murine CD11c MicroBeads UltraPure kit (Miltenyi Biotec, China) according to the manufacturer’s instructions.

### Cytokine assays

Cytokine levels in the serum and supernatant were quantified using enzyme-linked immunosorbent assay (ELISA) kits obtained from R&D Systems (USA) following the manufacturer’s instructions. Clec18a levels were analyzed using an ELISA kit sourced from Cusabio, China. The presented ELISA results are representative of ≥3 independent experiments. Data shown are from one representative experiment from at least triplicate wells (n refers to replicate wells).

### Immunoblotting and immunoprecipitation

The cells were lysed using radioimmunoprecipitation assay (RIPA) buffer (Beyotime, China), and protein concentrations were determined using a BCA assay kit (Thermo, China). Total protein was separated via SDS‒PAGE and transferred onto polyvinylidene fluoride (PVDF) membranes (Merck, Germany), followed by blocking with 5% nonfat milk in phosphate-buffered saline with Tween (TBST) (pH 7.5). The membranes were immunoblotted with primary antibodies overnight at 4°C and incubated with horseradish peroxidase (HRP)-conjugated secondary antibodies (Cell Signaling Technology, USA). The antibodies and reagents used are listed in Table S3. The protein bands were detected using an enhanced chemiluminescence kit (Pierce, USA) and visualized using a ChemiDoc XRS+ system (Bio-Rad, China).

Western blot experiments were independently repeated at least three times using biological replicates defined as independent cell culture preparations from independent experiments performed on different days. The blots shown are representative of one independent experiment; quantitative analyses (presented below) were performed using biological replicates.

For the immunoprecipitation assay, the cells were lysed in buffer containing 20 mM PIPES (pH 6.8), 1% Triton X-100, 150 mM NaCl, 150 mM sucrose, 0.2% sodium deoxycholate, 500 μM EDTA, and protease inhibitors on ice for 5 minutes. The samples were then centrifuged, and the supernatants were diluted to a concentration of 2 μg/mL in dilution buffer consisting of 20 mM PIPES (pH 6.8), 1% Triton X-100, 150 mM NaCl, 150 mM sucrose, 2.5 mM MgCl_2_ and 2.5 mM MnCl_2_. Primary antibody-conjugated protein A beads (Sigma, USA) were incubated at 4°C for two hours and washed with dilution buffer. Immunoblot analysis was performed using the methods described above.

### siRNA-mediated knockdown

siRNA-mediated knockdown was performed in PMs or BMDMs. The cells were cultured in half of the total volume of FBS-free medium and transfected with 3 ng/ml siRNA or control vector using INTERFEREin (Invitrogen, USA) for 6 hours. The remaining half of the culture medium was subsequently added, and the cells were cultured for another 48 hours. Next, the cells were subjected to subsequent experimental procedures. The sequence information for the siRNAs is provided in Table S3.

### Chromatin immunoprecipitation (ChIP) assay

PMs (1 × 107) were cultured in 150-mm plates. After LPS (100 ng/ml) treatment for 0.5 h, the cells were fixed, and the nuclei were spun down and resuspended in 500 μl of buffer (50 mM Tris at pH 8, 0.1% sodium dodecyl sulfate, and 5 mM EDTA). Sonication was performed in a Scientz-IID ultrasonic homogenizer with 10 cycles of 8 s each. All ChIP‒PCR steps were conducted with an EZ‒ChIP kit (Millipore, USA)[71] with an overnight immunoprecipitation step performed with anti-phospho-c-JUN antibody (5 μg, 3070S, CST) or control rabbit IgG (5 μg, 3900, CST).

### In vitro Mincle binding assay

The experiments were conducted as previously reported.[72] Briefly, rClec18a (2.5, 5, or 10 ng/ml) (purified with with endotoxin levels <0.05 EU/μg) (Cusabio, China) or TDB (2.5, 5, or 10 μg/ml) was coated onto 96-well plates (100 μl/well) overnight at 4°C. The coated plates were then washed with TBS and blocked for 1 h at room temperature. Mincle-Fc fusion protein (Recombinant Mouse CLEC4E Fc, R&D Systems, USA) and IgG1-Fc (Fc gamma R2B (FCGR2B), Invitrogen, USA) were incubated in plates with binding buffer for 2 hours at room temperature, followed by the addition of HRP-coupled goat anti-mouse IgG-Fc (Bethyl Laboratories, Texas) and the chromogenic substrate TMB (KPL laboratories). The plates were then analyzed using a microplate reader (TECAN, China) at 450 nm. To conduct the TDB-Clec18a competitive binding assay, the Mincle-Fc protein was incubated with rClec18a (10 ng/m) or TDB (10 μg/ml) for 10 minutes at room temperature before being added to with TDB- or rClec18a-bound plates, respectively. The mixture was then analyzed as described above. To conduct the Clec18a-coating assay and analyze Mincle endocytosis, rClec18a was coated onto plates as mentioned above, and PMs were then cultured and stimulated with LPS for 6 hours. Soluble rClec18a was added to a plate with LPS as a control.

### Surface plasmon resonance (SPR)

SPR measurements were performed on a Biacore instrument, and binding data were processed using the accompanying evaluation software. A carboxymethyl dextran sensor chip (CM5) was mounted in the instrument with the label side facing up. For ligand immobilization, the chip surface in flow cell 2 (FC2) was activated by standard amine-coupling chemistry using a 1:1 mixture of EDC and NHS. Recombinant Clec18a (rClec18a) or Mincle-Fc (as indicated below) was diluted in sodium acetate buffer and injected over the activated FC2 until the desired immobilization level was reached; flow cell 1 (FC1) was left unmodified as the reference channel. Remaining active esters were blocked according to the manufacturer’s amine-coupling protocol.

For the interaction analysis shown in Fig. C, rClec18a was immobilized on FC2 and Mincle-Fc was injected as the analyte in a concentration series (1, 0.5, 0.25, 0.125, 0.0625, and 0.03125 µM) over both FC1 and FC2. Each cycle consisted of an association phase followed by a dissociation phase, and the surface was regenerated between injections using an acidic regeneration solution (e.g., glycine-HCl, low pH) to restore the baseline. For the reverse configuration in Fig. D, Mincle-Fc was immobilized on FC2 and rClec18a was injected as the analyte using the same concentration series (1–0.03125 µM) and running scheme. Sensorgrams were double-referenced by subtracting signals from the reference channel (FC1) and blank injections. Kinetic parameters and/or binding affinity were obtained by global fitting to a 1:1 Langmuir binding model.

### qPCR

Total RNA was extracted using the TRIZOL reagent (Thermo Fisher, USA). A 10 μL reaction mixture containing 1 μg of total RNA, oligo (dT) primers, and a reverse transcription premix (Takara, Japan) was prepared and cDNA synthesis was subsequently performed. qPCR analysis was conducted using SYBR Green PCR Master Mix (Takara, Japan) on an ABI 7500 thermal cycler (Thermo Fisher, USA) under the following cycling conditions: initial denaturation at 95 °C for 3 min, followed by 40 cycles of denaturation at 95 °C for 10 s, annealing at 60 °C for 5 s, and extension at 72 °C for 10 s. mRNA levels were normalized to B2M expression as an internal control. The primer sequences are provided in Table S3.

### DNA transfection

The overexpression plasmids were obtained from Obio, China. RAW264.7 cells were cultured in DMEM supplemented with 10% fetal bovine serum (FBS). Before transfection, 4 μg of plasmid was mixed with the transfection reagent jetPEI (Polyplus, France). The resulting mixture was then added to the culture medium for a 24-hour transfection period. After 24 hours, the medium was replaced, and the cells were subjected to subsequent experiments.

### Exogenous mRNA transfection

Murin primary peritoneal macrophage were transfected at ∼60–80% confluency using jetMESSENGER (Polyplus, Shanghai, China), following the manufacturer’s guidance for mRNA (2.5μg mRNA: 5μl jetMESSENGER reagent) delivery in 6-well plates. In vitro–transcribed mRNA was synthesized and purified by Synbio Technologies (Suzhou, China), and the sequence were provided in the Supplementary Table S3. Purified mRNA was aliquoted in RNase-free tubes and stored at −80 l use. All procedures were performed using RNase-free consumables. Briefly, cells were seeded at a density suitable for 6-well plates in 2 mL complete medium per well. For each well of a 6-well plate, mRNA was diluted in 200 µL of the supplied jetMESSENGER mRNA buffer, mixed, and briefly centrifuged. jetMESSENGER reagent was vortexed briefly and added to the diluted mRNA, mixed gently, and incubated for 10–15 min at room temperature to allow complex formation. The transfection complexes were then added dropwise to cells in standard growth medium, followed by gentle rocking to distribute complexes evenly. After 24 hours of transfection, the cells were subjected to the following experiments.

### Whole-Cell Patch-Clamp Electrophysiology

Whole-cell patch-clamp recordings were performed at room temperature (20–25°C) using a MultiClamp 700B amplifier (Molecular Devices, USA). The recording setup was mounted on an upright infrared-differential interference contrast (IR-DIC) microscope (BX51WI, Olympus, Japan) equipped with a motorized micromanipulator (MP-325, Sutter Instrument, USA). Patch pipettes were pulled from borosilicate glass capillaries (O.D. 1.50 mm, I.D. 0.89 mm; VitalSense Scientific Instruments, China) using a P-97 micropipette puller (Sutter Instrument, USA) and polished with a microforge (MF-830, Narishige, Japan). Pipette resistance was maintained between 4 and 6 MΩ when filled with the internal solution.

To isolate TRPM7 currents and eliminate contamination from potassium channels, a cesium-based internal solution was referred to a previous published studies[44, 45] containing (in mM): 110 Cs-gluconate, 0.5 NaCl, 0.75 CaCl2, 10 HEPES, 10 HEDTA, 1.8 Cs4-BAPTA, and 2 Na2ATP (pH 7.3 adjusted with CsOH, ∼273 mOsm/kg). The extracellular solution contained (in mM): 135 Na-methanesulfonate, 5 Cs-gluconate, 2.5 CaCl2, 10 HEPES (pH 7.3 adjusted with NaOH, ∼280–290 mOsm/kg).

Data acquisition and stimulation protocols were controlled by pCLAMP 10.7 software (Molecular Devices, USA). TRPM7 currents were elicited using a voltage ramp protocol spanning from −100 mV to +150 mV over a duration of 250 ms, applied from a holding potential of −100 mV.

For experimental treatments, macrophages were stimulated with LPS (200 ng/mL) or LPS/rClec18a (20ng/ml). In pharmacological inhibition experiments, cells were pre-incubated with the TRPM7 channel inhibitor FTY720 (10 µM) or the kinase inhibitor TG100-115 (10 µM) for 0.5 hour prior to LPS stimulation. The inhibitors were maintained in the bath solution throughout the subsequent recording period to ensure continuous inhibition.

### In vitro ubiquitination assay

The E3 ubiquitin ligase activity assay was performed following a previously described method.[39] Briefly, the reaction mixture was prepared with 100 nM E1, 0.5 mM UbcH7 (E2), 5 μM ubiquitin, and 2 μM ATP in ULR buffer (25 mM Tris-HCl, pH 8.0; 100 mM NaCl; 2.0 mM MgCl2; and 1 mM dithiothreitol). Cbl was immunoprecipitated from Cbl-overexpressing HEK293 cells and then added to the mixture. In the activation assay, EGTA-buffered CaCl2 solution or the HEK293 cell homogenate fraction (5 μl/each sample) was added to the immunoprecipitated Cbl and then added to the reaction mixture. The reaction was carried out at 37 °C for 3 h and stopped by the addition of 25 μl of 2×SDS‒PAGE sample buffer. The reaction mixtures were then subjected to immunoblotting.

### Flow cytometry

Single-cell BMDM or PM suspensions were washed, blocked with a 2.4G2 antibody (Bio X cell), and stained with the appropriate antibodies in MACS buffer (PBS+1% FBS) to detect Mincle. Fluorescence data for 10^5^ events were acquired on a FACS LSR II (BD Bioscience, USA) and analyzed using FlowJo software (Tomy Digital Biology Co., Ltd., Japan).

### Pull-down assays

For the GST or HIS pull-down assays, equal amounts (0.5 mg) of the GST-tagged fusion protein or HIS-tagged fusion protein were briefly mixed and incubated on ice for 3 h. Subsequently, the mixture was loaded onto Glutathione Sepharose 4 B or Ni-agarose resin columns. After washing five times with wash buffer, the proteins were eluted with wash buffer supplemented with 15 mM reduced glutathione. The eluates were separated using 12% SDS‒PAGE, transferred to PVDF membranes (Millipore, Billerica, MA, USA), and probed with anti-HIS (or anti-GST) antibodies (Sigma‒Aldrich, Merck KGaA, Darmstadt, Germany). GST and HIS from Wuhan Genecreate (Wuhan, China) were used as negative controls. Three replicates were used for each pull-down assay.

### Immunofluorescence and calcium imaging

For immunofluorescence staining of Clec18a, murine BMDMs were plated on slides and fixed with 4% paraformaldehyde (PFA). The cells were then washed with PBS, permeabilized with a membrane-disrupting solution (50–100 μl), and incubated at room temperature for 20 minutes. The cells were subsequently washed three times with PBS for 5 minutes each. Next, the cells were blocked with 3% bovine serum albumin (BSA) at room temperature for 30 min. Next, the cells were incubated overnight at 4°C with an anti-mouse Clec18a antibody. After three washes, the membranes were incubated with secondary antibodies at room temperature for 50 min. The slides were rinsed three times in PBS (pH 7.4) by shaking on a decolorizing shaker. DAPI dye was then applied to the samples, which were subsequently incubated for 10 min at room temperature in the dark. After decolorization, the slides were gently dried and sealed with an antifluorescence quenching sealant before being subjected to confocal microscopy analysis using a Nikon Eclipse Ti & C2 system. The excitation wavelength of DAPI ranged from 330 nm to 380 nm with an emission wavelength of 420 nm, and the excitation wavelength of FITC ranged from 465 nm to 495 nm, while its emission wavelength ranged from 515 nm to 555 nm.

For calcium imaging, BMDMs previously plated on glass coverslips were loaded with the calcium indicator Fluo-4-AM (5 μM, Invitrogen, Thermo Fisher Scientific) for 30 minutes at 37°C and then resuspended in normal external buffer (140 mM NaCl, 5 mM KCl, 2 mM CaCl2, 2 mM MgCl_2_, and 10 mM HEPES, titrated to pH 7.4 with NaOH). Live calcium imaging was performed in imaging buffer (Hank’s balanced salt solution supplemented with 2 mM Ca2+, Lonza) using a Leica Stellaris 5 laser scanning confocal microscope. Measurements were performed by continuous perfusion of the bath solution with a gravity feed system, and ionomycin (1 μM) was perfused as a positive control for cellular responsiveness. The experiments were performed in normal external buffer supplemented with 500 ng/ml LPS, as indicated in the figures and legends. Images of macrophages at an excitation wavelength of 488 nm (0.5% power) were captured every 5 s for the indicated time of 5 min. Regions of interest (ROIs) were drawn on one distinct cell per field; Fluo-4 traces were generated by averaging the pixel signals within the ROIs and were normalized to the baseline fluorescence intensity of the first five frames (ΔF/F0). Control experiments were performed using murine macrophages loaded with the calcium indicator alone. For the analysis of calcium signaling events, one ROI per cell was designed, and single-cell Fluo-4 traces were generated by averaging pixel signals within the ROIs and were normalized to the baseline fluorescence intensity of the first five frames (ΔF/F0) using ImageJ software.

### Statistical analysis

The obtained results were subjected to statistical analysis using Student’s t test or analysis of variance (one-way or two-way ANOVA) with appropriate multiple comparisons, as applicable, using Prism software (version 10.0; GraphPad Software, Inc.). Animals were randomly allocated to experimental groups, and outcome assessments were performed by investigators blinded to group assignment whenever feasible. Kaplan‒Meier survival curves were compared using the log-rank Mantel‒Cox test. All tests were two-tailed unless otherwise specified. A *p* value less than 0.05 was considered statistically significant. Data are presented as mean ± SD (or mean ± SEM, as indicated in figure legends). Biological replicates denote the individual animals number indicated in the experiments for in vivo assay. Replicate wells denote the technical replicates of in vitro study.

## DATA AVAILABILITY STATEMENT

All data supporting the findings of this study are available within the article and its supplementary information. Uncropped Western blot and flow cytometry data are provided in Raw data file.

## AUTHOR CONTRIBUTIONS

Zhengyu Jiang: Conceptualization, Formal analysis, Investigation, Writing – original draft.

Na Li: Investigation.

Huabiao Chen: Investigation.

Lulong Bo: Investigation.

Tao Li: Investigation.

Yan Zhang: Conceptualization, Formal analysis, Writing – original draft, Supervision.

Xiaoming Deng: Conceptualization, Formal analysis, Writing – original draft, Supervision.

Jinjun Bian: Conceptualization, Formal analysis, Writing – original draft, Supervision, Funding acquisition.

## ACKNOWLEDGEMENTS

This study is supported by the National Natural Science Foundation of China (81771697, 81772105 82000560, 82272205), Shanghai Key Laboratory of Nautical Medicine and Translation of Drugs and Medical Devices, Committee of Science and Technology of Changning District, Shanghai (CNKW2022Y59), and Research program of Changhai Hospital (2023PY32).

## ETHICS AND CLINICAL TRIAL REGISTRATION

Clinical sample research in our work is approved by the Ethics Committee of Changhai Hospital, Naval Medical University (Approval No. CHEC2018-164) and registration with the China Clinical Trials Registry (ChiCTR1800015714). Written consent was obtained from patients enrolled. Ethical approval for the animal experiments was obtained from the Ethical Committee of Changhai Hospital following the Experimental Animal Regulations of Naval Medical University.

## LIST OF NON-STANDARD ABBREVIATIONS

Act D: actinomycin D;
BMDCs: bone marrow-derived dendritic cells;
BMDMs: bone marrow-derived macrophages;
CHX: cycloheximide;
CLP: cecal ligation and puncture;
Cbl: Casitas B-lineage lymphoma;
Clec18a: C-type lectin 18A;
FIP: fecal-induced peritonitis;
Mincle: macrophage-inducible C-type lectin;
PMs: peritoneal macrophages;
rClec18a: recombinant Clec18a;
SPR: surface plasmon resonance;
TDB: trehalose-6,6-dibehenate;
Trpm7: transient receptor potential melastatin-like 7.

## EXPANDED VIEW

Figures S1–S5 and Tables S1–S3

## SUPPLEMENTARY DATA DESCRIPTION

Supplementary material includes five supplementary figures (Figures S1–S5) demonstrating Clec18a expression patterns, functional characterization, Mincle receptor binding and endocytosis, Cbl-mediated ubiquitination mechanisms, and Trpm7-calcium signaling; three supplementary tables (Tables S1–S3) listing clinical cohort demographics, GEO dataset details, and experimental reagent and sequences; together with raw data files containing uncropped Western blot images and original flow cytometry data (Raw data file).

## SUPPLEMENTARY MATERIALS

### Supplementary Figure legends

**Figure S1.**
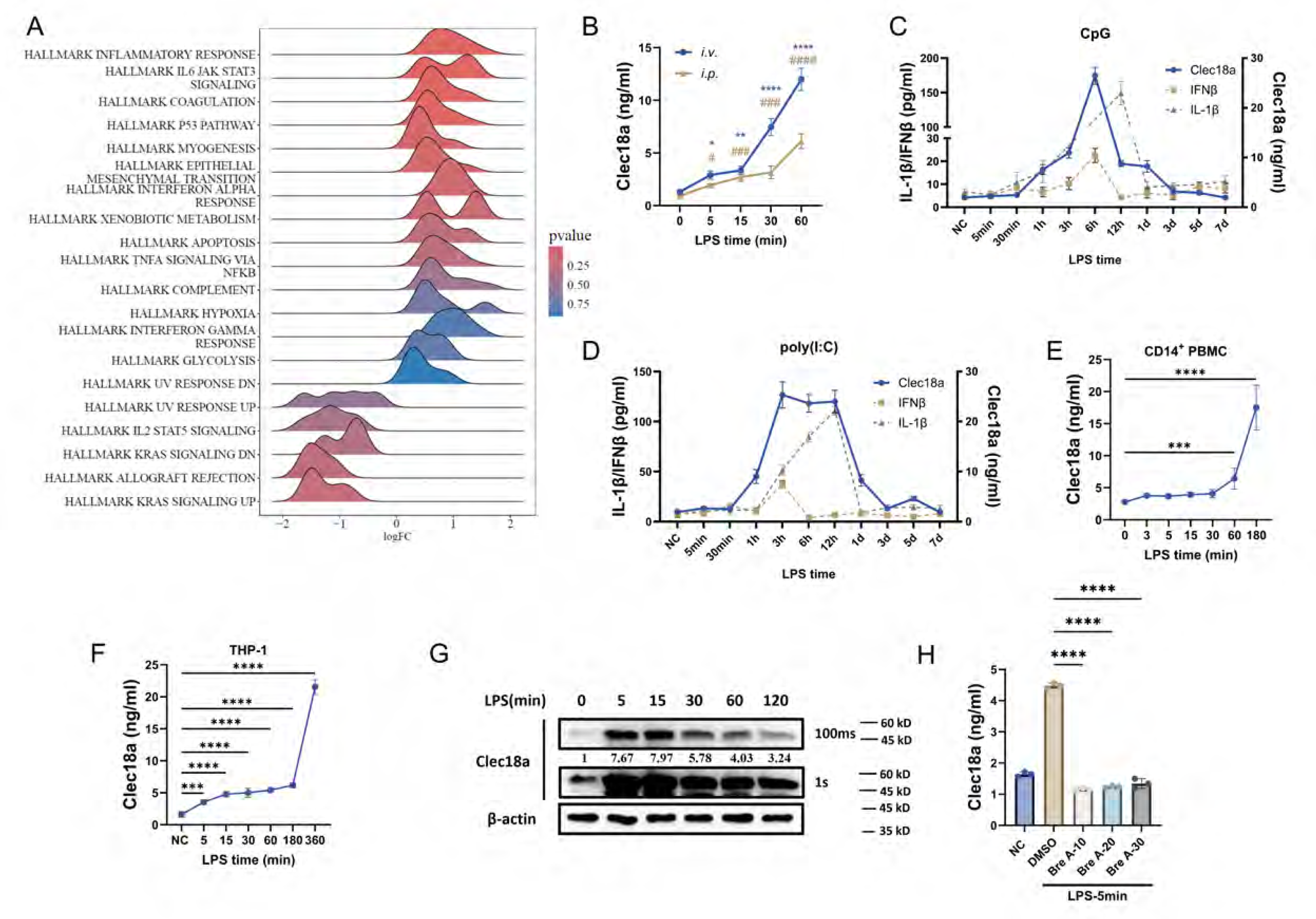
Expression pattern of Clec18a and its relationship with inflammatory cytokines. A: Gene set enrichment analysis (GSEA) of Clec18a-high samples from merged sepsis dataset. Vertical axis represents the normalized enrichment score (NES). LogFC>0 reflects the up-regulation of the indicated pathway. B: Levels of Clec18a in the serum of mice injected intravenously (i.v.) or intraperitoneally (i.p.) with LPS (10 mg/kg) at various time points (n=4 per group per timepoints). C, D: Cytokine and Clec18a levels in the serum of C57BL/6 mice injected intravenously with CpG (C) (50μg/mouse) or poly(I:C) (50μg/mouse) (D) at the indicated time points (n=5 per group per timepoints). E, F: Levels of Clec18a in the supernatant of human CD14^+^ PBMCs (E), PMA-stimulated THP-1 cells (F), and stimulated with LPS (10 ng/ml) at various time points (n=4). G: Western blot analysis of Clec18a in BMDMs stimulated with LPS at various time points, with two lanes of the same blot showing the results of the different LPS exposure times (100 ms and 1 s). H: Clec18a level in the supernatant of BMDMs pre-treated with Brefeldin A (10, 20 and 30mg/ml) for 5 hours and then stimulated with LPS for five minutes (n=3). *p<0.05, **p<0.01, ***p<0.001, ****p<0.0001; one-way ANOVA (B, E, F, H); n=biological replicates for in vivo assay and replicate wells for in vitro assay; Data shown in all panels are representative of at least three independent experiments.

**Figure S2.**
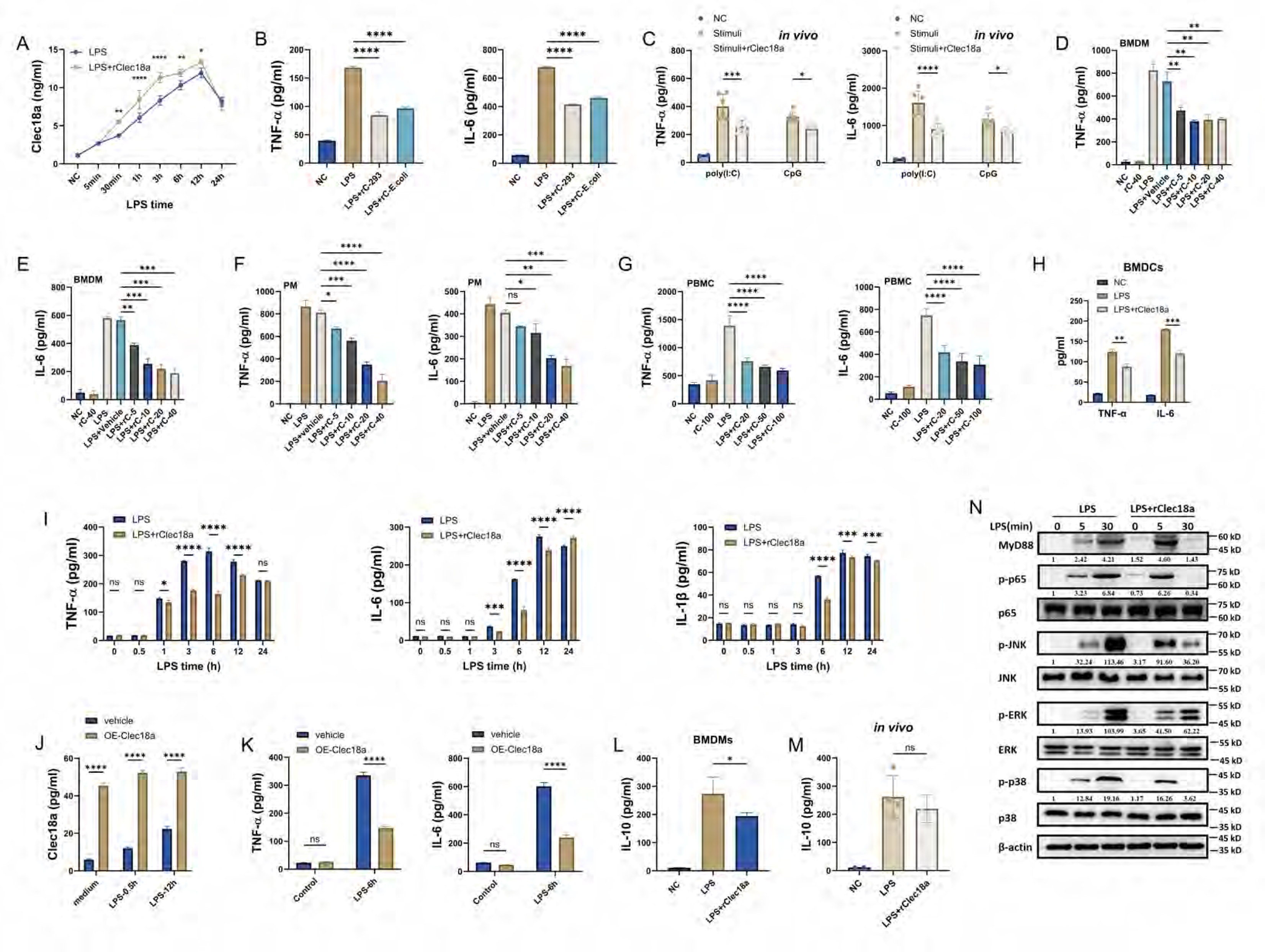
Clec18a alleviates inflammation. A: Clec18a level in the serum of C57BL/6J mice injected with LPS or LPS+rClec18a *i.p.* (20ng/g) for indicated timepoints (n=4 per group per timepoints). B: TNF-α and IL-6 levels in the supernatants of BMDMs treated with 293T-derived rClec18a (rC-293) and *E. coli.*-derived rClec18a (rC-E.coli) and stimulated with LPS for 6 hours (n=3). C: TNF-α and IL-6 levels in the serum of C57BL/6 mice treated with CpG (50 µg/mouse) or poly(I:C) (50 µg/mouse) with rCelc18a (20 ng/g) (stimuli indicates CpG or poly(I:C)) for 6 hours (n=6 per group). D-H: TNF-α and IL-6 levels in the supernatants of BMDMs (D, E), PMs (F), human PBMCs (G) and murine BMDCs (H) treated simultaneously with human or mouse rClec18a at indicated concentrations (ng/ml) (rClec18a at 20ng/ml for BMDCs) and LPS for 6 hours (n=3). I: TNF-α, IL-6 and IL-1β levels in the supernatants of BMDMs treated LPS and LPS+rClec18a for indicated timepoints (n=3). J, K: Levels of Clec18a (J), TNF-α, and IL-6 (K) in the supernatants of RAW264.7 cells transfected with the Clec18a overexpression plasmid (OE-Clec18a) or control vector for 24 hours and then stimulated with LPS at various time points (n=3). L, M: IL-10 levels in the supernatant of PMs (L) and the serum of mice (M) treated with LPS or LPS+rClec18a for 6 hours (n=4 per group). N: Western blot analysis of PMs stimulated with LPS or LPS+rClec18a at various time points. *p<0.05, **p<0.01, ***p<0.001, ****p<0.0001, ns: no significance; one-way ANOVA (B-H, L-M); two-way ANOVA (I-K); n=biological replicates for in vivo assay and replicate wells for in vitro assay; Data shown in all panels are representative of at least three independent experiments.

**Figure S3.**
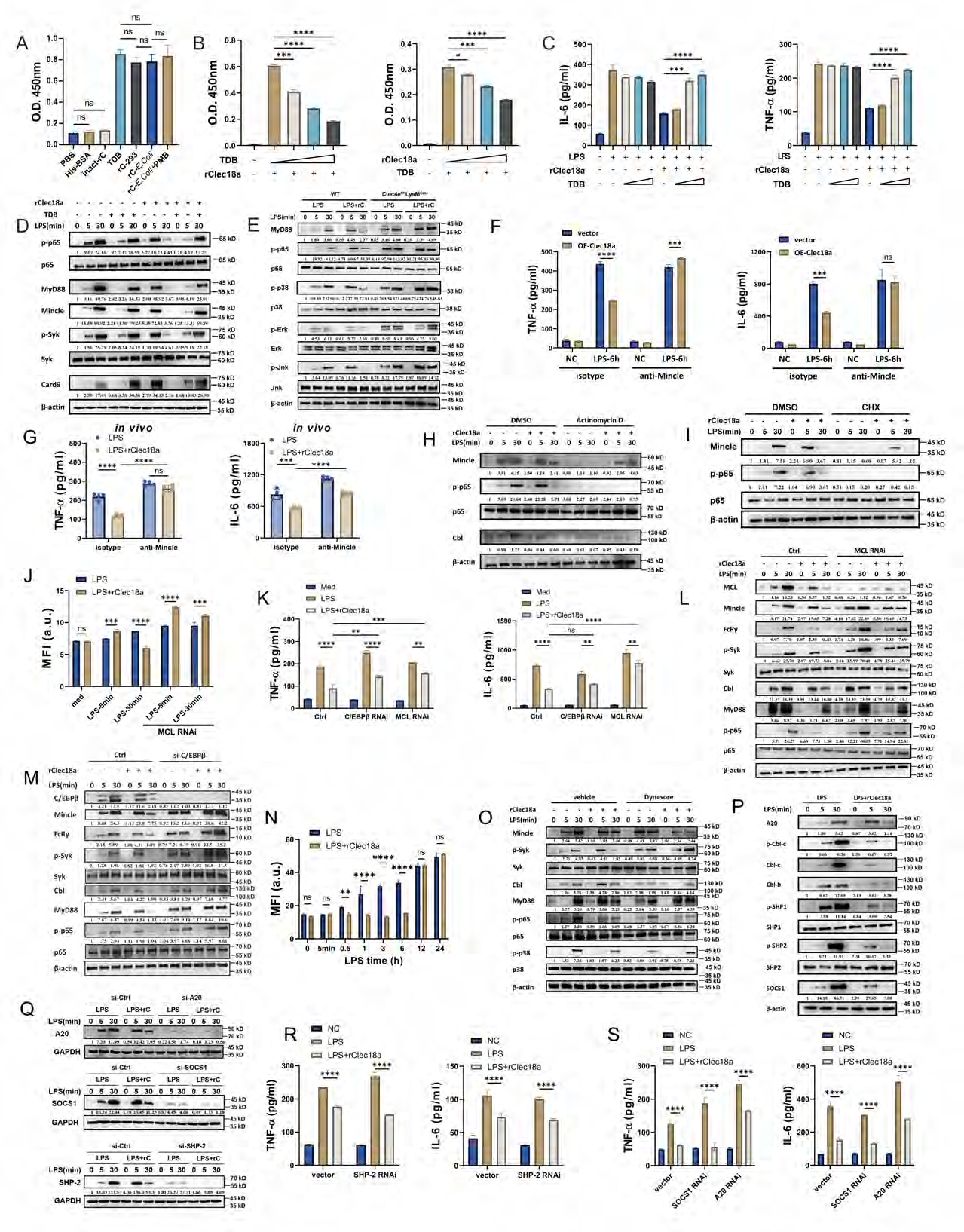
Clec18a regulates TLR4 signaling via its receptor, Mincle. A: Cell-free binding assay (O.D. 450nm) of Mincle-Fc with His-tagged BSA, heat-inactivated rClec18a (derived from *E. coli*, 10ng/ml), 283T cell-derived rClec18a (rC-293, 10ng/ml), *E. coli.*-derived rClec18a (rC-*E. coli.*, 10ng/ml) or rC-*E. coli* cotreated with polymyxin B (PMB, 10μg/ml) (n=3). B: TDB-Clec18a competitive binding assay of Mincle-Fc protein which was incubated with rClec18a (10 ng/m) or TDB (10 μg/ml) before being added to with TDB-(2.5, 5 or 10 μg/ml) or rClec18a-(2.5, 5 or 10 μg/ml)bound plates, respectively. C: TNF-α and IL-6 levels in the supernatant of BMDMs treated with rClec18a (20 ng/ml) in TDB (2.5, 5, or 10 μg/ml) bound plates and stimulated with LPS for 6 hours (n=3). D: Western blot analysis of BMDMs treated with LPS and rClec18a or TDB for the indicated time points. E: Western blot analysis of BMDMs from Clec4e^fl/fl^LysM^Cre+^ and littermate wild-type (WT) mice treated with LPS and LPS+rC for indicated timepoints. F: TNF-α and IL-6 levels in the supernatant of RAW264.7 cells with transfected with Clec18a overexpression (OE-Clec18a) plasmid and treated with Mincle antibody (10 μg/mouse) or isotype control (10 μg/mouse) for 30 minutes and then with LPS for 6 hours (n=3). G: TNF-α and IL-6 levels in the serum of mice treated with an anti-Mincle antibody or isotype control and then stimulated with LPS or LPS+rClec18a for 6 hours (n=4 per group). H, I: Western blot analysis of BMDMs pretreated with Act D (10 μM) (H) or CHX (30 μM) (I) for 30 minutes and then stimulated with LPS for various durations. J: Flow cytometry analysis of Mincle mean fluorescence intensity (MFI) in BMDMs with MCL siRNA knockdown and treated with the indicated reagents (n=3). K-M: TNF-α and IL-6 levels in the supernatant (K) and Western blot analysis (L, M) of PMs with MCL or C/EBPβ siRNA knockdown and treated with LPS or LPS+rClec18a for 6 hours (K) or various time points (L, M) (n=3). N: Flow cytometry analysis of Mincle mean fluorescence intensity (MFI) in BMDMs treated with LPS or LPS+rClec18a at the indicated timepoints (n=3). O: Western blot analysis of PMs pre-treated with Dynasore for 30min and then stimulated with LPS or LPS+rClec18a at indicated timepoints. P: Western blot analysis of PMs treated with LPS or LPS+rClec18a for various durations (n=3). Q-S: Western blot of knock down efficiency of targeted genes (Q) and TNF-α and IL-6 levels (R and S) in the supernatant of PMs with SOCS1, A20 and SHP-2 knockdown and treated with LPS or LPS+rClec18a for 6 hours. *p<0.05, **p<0.01, ***p<0.001, ****p<0.0001; one-way ANOVA (A-C), two-way ANOVA (F, G, J, K, N, R, S); n=biological replicates for in vivo assay and replicate wells for in vitro assay; Data shown in all panels are representative of at least three independent experiments.

**Figure S4.**
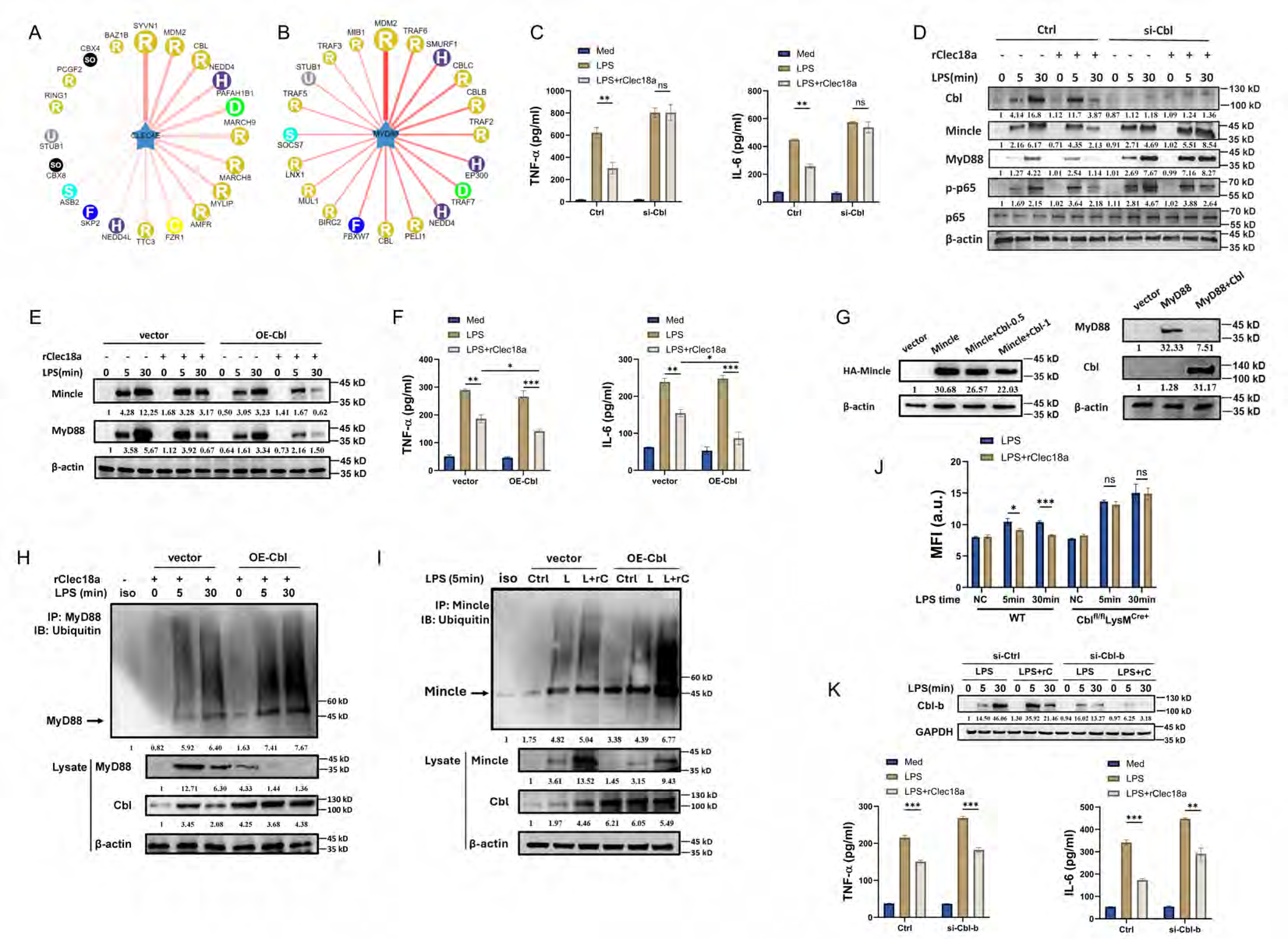
Clec18a induces Mincle and MyD88 degradation through Cbl-mediated ubiquitination. A, B: Illustration of predicted ubiquitin E3 ligases interacting with Mincle (Clec4e) and MyD88, retrieved from http://ubibrowser.bio-it.cn/ubibrowser/. C, D: TNF-α and IL-6 levels (C) in the supernatant and western blot analysis (D) of PMs with Cbl siRNA knockdown for 48 hours of PMs and treated with LPS and rClec18a (n=3). E-F: Western blot analysis Cbl-overexpressed RAW264.7 cells (E) simulated with LPS and rClec18a at indicated time points and with TNF-α and IL-6 levels in the supernatant (F) at 6 hours stimulated. G: Western blot analysis of HA-Mincle in RAW264.7 cells co-transfected with overexpression plasmids of Mincle and Cbl, or MyD88 and Cbl, Cbl-0.5 and Cbl-1 represent plasmids of (0.5 or 1 μg/ml). H, I: Western blot analysis of ubiquitin of immunoprecipitated Mincle (H) and MyD88 (I) in Cbl-overexpressed RAW264.7 cells treated with LPS and rClec18a for indicated time points. Lysate from LPS-5min (vector) and LPS-30min (vector) used in isotype lane. J: Flow cytometry analysis of Mincle on the cell surface in BMDMs of WT and Cbl^fl/fl^LysMCre^+^ mice and treated with LPS or LPS+rClec18a for indicated time points (n=3). K: Western blot of knock down efficiency of Cbl-b and TNF-α and IL-6 levels in the supernatant of PMs with Cbl-b siRNA for 48 hours and treated with LPS and rClec18a for 6 hours (n=3). *p<0.05, **p<0.01, ***p<0.001, ****p<0.0001, ns: not significant; two-way ANOVA; n=biological replicates for in vivo assay and replicate wells for in vitro assay; Data shown in all panels are representative of at least three independent experiments.

**Figure S5.**
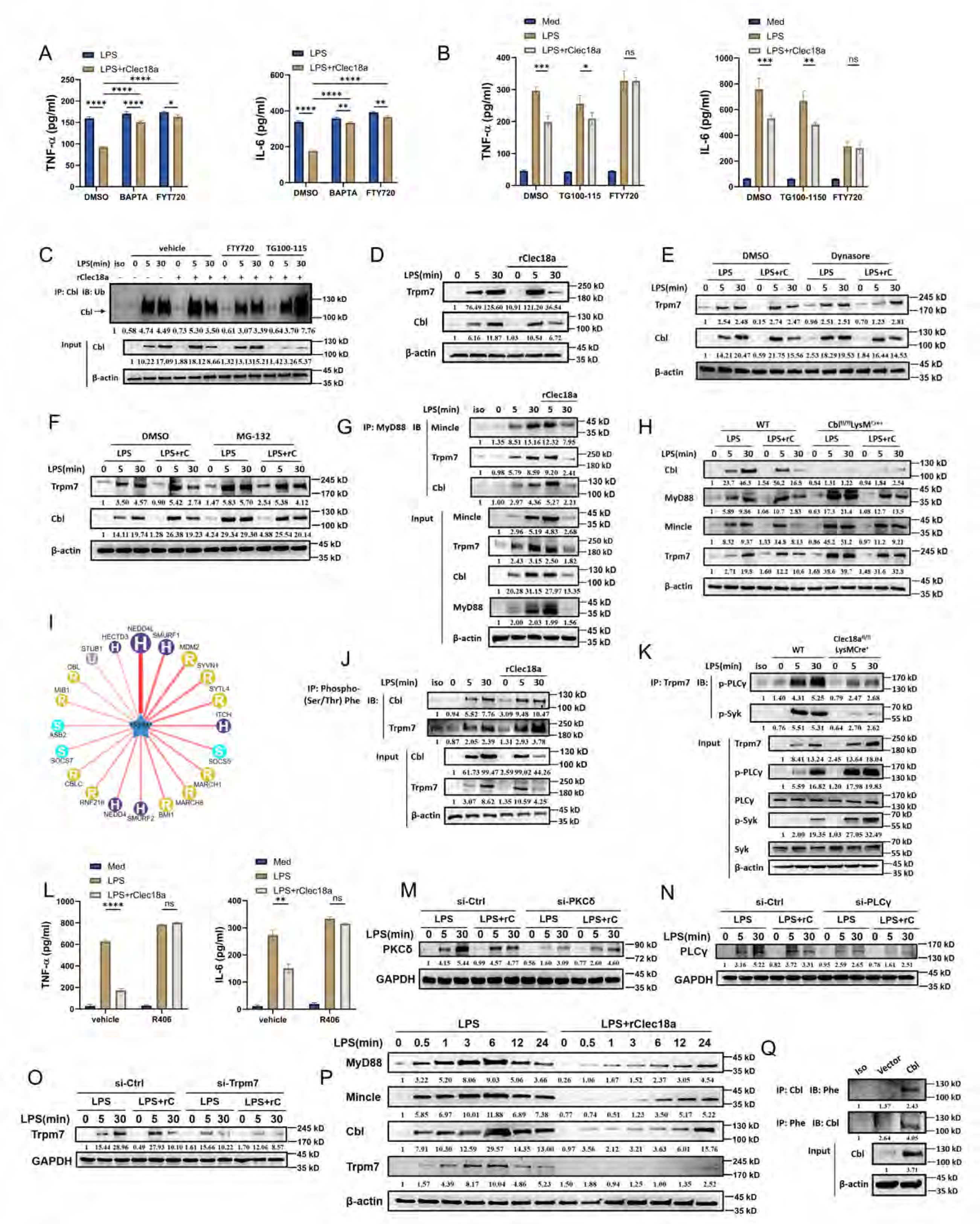
Trpm7-derived calcium is responsible for Clec18a-mediated Cbl activation. A, B: TNF-α and IL-6 levels in the supernatant of PMs pretreated with BAPTA (calcium chelator, 22 μM), FTY720 (Trpm7 channel inhibitor, 100 μM), or TG100-150 (Trpm7 kinase inhibitor, 10 μM) for 30 minutes and then stimulated with LPS or LPS+rClec18a for 6 hours (n=3). C: Ubiquitination assay of immunoprecipitated Cbl in PMs pretreated with FTY720 or TG100-150 and stimulated with LPS or LPS+rClec18a for the indicated time points. D: Western blot analysis of BMDMs treated with LPS and rClec18a for the indicated durations. E, F: Western blot analysis of BMDMs pretreated with Dynasore (10 μM) (E) or MG-132 (20 μM) (F) and then treated with LPS and rClec18a for the indicated durations. G: Immunoprecipitation of MyD88 with Mincle, Trpm7, and Cbl in PMs treated with LPS or LPS+rClec18a for the indicated durations. Lysate from LPS-30min (vehicle) used in isotype lane. H: Western blot analysis of BMDMs from WT or Cbl^fl/fl^LysMCre^+^mice treated with LPS or LPS+rClec18a at the indicated time points. I: Illustration of predicted ubiquitin E3 ligases interacting with Trpm7, retrieved from http://ubibrowser.bio-it.cn/ubibrowser/. J: Immunoprecipitation of phosphor-(Ser/Thr) Phe from Cbl and Trpm7 in PMs treated with LPS and rClec18a for the indicated durations. K: Immunoprecipitation of Trpm7 with p-PLCγ and p-Syk in PMs from WT or Clec18a^fl/fl^LysMCre^+^ mice treated with LPS for the indicated durations. Lysate from LPS-30min (WT) used in isotype lane. L: TNF-α and IL-6 levels in the supernatant of PMs pretreated with R406 and stimulated with LPS or LPS+rClec18a for 6 hours (n=3). M-O: Western blot of knock down efficiency of targeted genes of PMs with siRNA for 48 hours and stimulated with LPS or LPS+rClec18a for indicated time points. P: Western blot analysis of BMDMs stimulated with LPS and rClec18a for the indicated durations. Q: Immunoprecipitation of phosphor-(Ser/Thr) Phe from Cbl and vice versa in RAW264.7 cells overexpressing Cbl. Lysate from Cbl transfection used in isotype lane; n=biological replicates for in vivo assay and replicate wells for in vitro assay; Data shown in all panels are representative of at least three independent experiments.*p<0.05, **p<0.01, ***p<0.001, ****p<0.0001, ns: not significant; two-way ANOVA.

### Supplementary TABLES

**Supplementary Table S1. CLEC18A_expression_across_4GSEs.csv**

**Supplementary Table S2. The baseline of 21 subjects included in the study.**

**Supplementary Table S3. Reagents and sequence.**

